# Development of a multi-sensor integrated midbrain organoid-on-a-chip platform for studying Parkinson’s disease

**DOI:** 10.1101/2022.08.19.504522

**Authors:** Sarah Spitz, Silvia Bolognin, Konstanze Brandauer, Julia Füßl, Patrick Schuller, Silvia Schobesberger, Christian Jordan, Barbara Schädl, Johannes Grillari, Heinz D. Wanzenboeck, Torsten Mayr, Michael Harasek, Jens C. Schwamborn, Peter Ertl

## Abstract

Due to its ability to recapitulate key pathological processes *in vitro*, midbrain organoid technology has significantly advanced the modeling of Parkinson’s disease over the last few years. However, some limitations such as insufficient tissue differentiation and maturation, deficient nutrient supply, and low analytical accessibility persist, altogether restricting the technology from reaching its full potential. To overcome these drawbacks, we have developed a multi-sensor integrated organ-on-a-chip platform capable of monitoring the electrophysiological, respiratory, and dopaminergic activity of human midbrain organoids. Our study showed that microfluidic cultivation resulted in a marked reduction in necrotic core formation, improved tissue differentiation as well as the recapitulation of key pathological hallmarks. Non-invasive monitoring employing an orthogonal sensing strategy revealed a clear time dependency in the onset of Parkinson’s disease-related phenotypes, reflecting the complex progression of the neurodegenerative disorder. Furthermore, drug-mediated rescue effects were observed after treatment with the repurposed compound 2-hydroxypropyl β-cyclodextrin, highlighting the platform’s potential in the context of drug screening applications as well as personalized medicine.

## 1. Introduction

With a prevalence of 9.4 million, Parkinson’s disease (PD) constitutes the second most common neurodegenerative disorder worldwide.^1^ Characterized by intracellular inclusions of α-synuclein and the loss of dopaminergic neurons within the *substantia nigra* of the human midbrain, this heterogeneous disease results in a variety of debilitating motor and non-motor symptoms.^2^ Whereas the lack of disease-modifying and neuroprotective strategies has restricted the treatment of PD to symptomatic control, the development of new therapeutics remains hampered by the high failure rates observed in clinical trials. This ill success of putative drug candidates can be attributed, at least in part, to the inability of current disease models to replicate critical pathological hallmarks of the disease.^3^ With the emergence of induced pluripotent stem cell (iPSC)-technology, however, new possibilities toward physiologically relevant and patient-specific *in vitro* models have opened up in the form of so-called “organoids”.^4,5^ Organoids denote *in vitro* derived microtissues that, by undergoing some level of self-organization, can recapitulate fundamental physiological facets of *in vivo* organs. Several organoid-based models of the human brain have been developed so far, including microtissues of the forebrain, the hindbrain, and of the midbrain, the afflicted region in PD.^6–8^ Midbrain organoid models not only exhibit essential features of the tissue’s three-dimensional (3D) cytoarchitectural arrangement and function but also mirror pathological hallmarks of PD including dopaminergic neurodegeneration.^9–12^ Despite these significant advancements, however, several challenges persist, including the lack of specific cell types (e.g., endothelial cells), the tissue’s immaturity, organoid variability, and nutrient deficiency-based growth restrictions – a result of the organoid’s inherent structure and the concomitant lack of vasculature. Moreover, organoid technology still strongly relies (a) on unphysiological cultivation conditions that omit critical biophysical cues and (b) on the use of invasive endpoint analyses that hinder the investigation of progressive developmental processes. One important biophysical cue that has been largely overlooked is the brain-specific interstitial fluid flow, which has been linked to a variety of essential functions, including the delivery of nutrients, the removal of metabolic and neurotoxic waste, non-synaptic cell-cell communication, tissue homeostasis as well as cell migration.^13,14^ Moreover, as part of the so-called “glymphatic system” – a glial-mediated clearance system of the human brain – interstitial fluid flow has been connected to exacerbated protein deposition in models of Alzheimer’s disease and PD as well as to the delivery and clearance of drugs, making it of considerable importance in the context of pharmacological screening applications.^13,15,16^

One technology capable of applying defined flow profiles, fluid velocities, and shear forces is microfluidics.^17^ Its ability to reproducibly control various parameters such as mechanical stimulation, laminar fluid dynamics (e.g., perfusion), temperature profiles, and gaseous permeability has already led to the successful recreation of various tissue models *in vitro* in the form of so-called organs-on-a-chip.^18–20^ Although organ-on-a-chip technology has expanded massively over the last decade, only a handful of studies have set out to combine brain organoids with organ-on-a-chip technology.^7,21–28^ For example, brain-on-a-chip platforms have already been successfully employed to assess the effects of prenatal exposure to various neuroteratogens such as cadmium, nicotine, and cannabis, to study the physical underpinnings of human brain folding as well as to model neural tube development *in vitro*.^7,22,25–27^ A more recent study by Cho *et al*.^24^ demonstrated significant improvements in the functional maturation of human brain organoids embedded within a brain extracellular matrix enriched hydrogel using gravity-driven flow. Next to improved cellular viabilities and reduced necrotic cores, dynamic cultivation could markedly reduce organoid size variation from a coefficient of variance of 43.4 % to 17.4%. However, none of the above-referenced studies have investigated neurodegenerative processes or exploited the potential of microfluidic technology to integrate in-line sensing strategies needed to increase analytical accessibility, a prerequisite of any high-quality *in vitro* tissue model.

Here, we present, for the first time, a multi-sensor integrated microfluidic platform for the dynamic long-term cultivation and analysis of human iPSC-derived midbrain organoids. The main design feature of the organoid-on-a-chip platform consists of unidirectional medium flow through three interconnected chambers, each equipped with an optical, electrical or electrochemical microsensor (Fig. 1). Dynamic cultivation of the organoids is accomplished by combining a hydrogel-based flow restrictor with gravity-driven flow to simulate interstitial flow profiles.^29^ To provide information on essential cellular parameters, including cell growth, metabolic activity, tissue viability, and differentiation, as well as cellular pathology, an optical, luminescent-based oxygen sensor spot is located in the organoid chamber.^30–32^ Additionally, a multi-electrode array is integrated to non-invasively monitor the electrophysiological activity of neuronal processes, an essential prerequisite for functional cell coupling and tissue maturation. Furthermore, an enzyme-based amperometric sensor is introduced into the downstream compartment to monitor the release of the neurotransmitter dopamine, as it directly reflects the maturation progress and pathological status of midbrain tissues. As this study sets out to investigate pathophysiological alterations in PD, midbrain organoids derived from a PD patient carrying a triplication mutation of the α-synuclein gene (3xSNCA) have been selected. Under physiological conditions, α-synuclein modulates crucial cellular processes such as synaptic vesicle trafficking, neurotransmitter release, and neuronal differentiation.^33,34^ Due to its structural instability, however, α-synuclein can adopt several conformations that favor oligomerization and aggregation, which have been associated with neurotoxicity and neurodegeneration.^35^ Elevated levels of α-synuclein (e.g., through multiplications of the gene) further reinforce the protein’s propensity to aggregate and have been associated with full PD penetrance.^36–38^ We have previously shown that the compound 2-hydroxypropyl β-cyclodextrin (HP-β-CD) can rescue PD-related phenotypes in a personalized PD model carrying biallelic pathogenic variants in the PINK1 gene by elevating the neuronal autophagy and mitophagy capacity.^39^ Autophagy dysregulation has been implicated in many genetic models of PD including that of α-synuclein, rendering HP-β-CD, an ideal candidate for the evaluation of the microfluidic platform.

**Figure 1:**
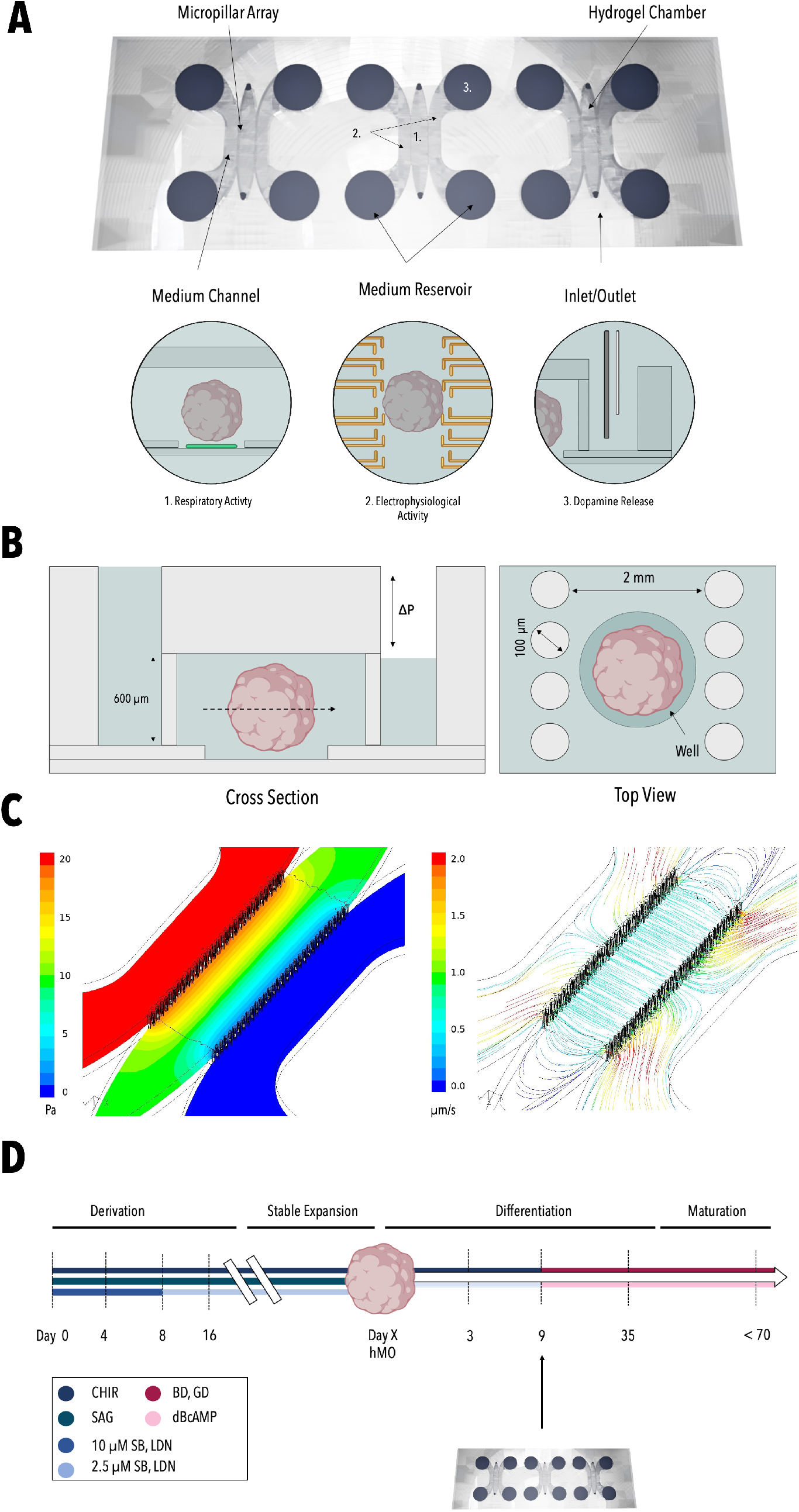
Render of the PDMS-based microfluidic device (Fusion 360) comprised of three individual culture chambers, including the positioning of 1. the optical oxygen sensor, 2. the multi-electrode array (MEA), and 3. the electrochemical dopamine sensor. (A). Cross view and top view of the microfluidic organ-on-a-chip platform (B). Results of the CFD simulation depicting the pressure profile on-chip (left panel) and the generated laminar flow profile (right panel) (C). Workflow of organoid maturation and microfluidic cultivation (D).

Thus, in our final set of experiments, the multi-sensor integrated platform is employed to investigate the dynamic exposure of HP-β-CD on PD onset and progression.

## 2. Results and Discussion

### 2.1 Hydrostatic pressure-driven flow emulates cerebral interstitial fluid flow regimes *in vitro*

To account for interstitial fluid flow in our sensor-integrated *in vitro* model, a combination of hydrostatic pressure-driven flow and hydrogel-based fluid restriction was explored.^29^ As depicted in Figure 1A, each cultivation chamber of the PDMS-based organ-on-a-chip platform is comprised of three individual compartments interconnected by two micropillar arrays. While the two outer chambers form the medium channels, the middle chamber is designed to accommodate the differentiating human midbrain organoid (hMO) in a three-dimensional matrix, emulating *in vivo* elasticities (Matrigel^®^).^40,41^ To facilitate the reproducible positioning of the organoids in the microfluidic device, a 250 μm deep cylindrical well was integrated at the bottom of the central hydrogel chamber (Fig. 1B). Hydrostatic pressure-driven flow is generated by filling up both reservoirs of the supply channel (left medium channel), which subsequently directs medium flow through the hydrogel matrix and the embedded microtissue within towards the waste compartment (right medium channel) (Fig. 1B). Therefore, nutrient supply is no longer restricted to diffusion, but nutrients are actively transported to the embedded organoid via convective flow. It is important to note that the integrated hydrogel-restrictor keeps shear forces at a minimum, thus fulfilling an essential requirement in the cultivation of neuronal microtissues. To gain a deeper understanding of the applied flow behavior within the microfluidic platform, a computational fluid dynamic (CFD) simulation was performed. In-silico analysis demonstrated that the main pressure drop occurs at the hydrogel interface (Fig. 1C), resulting in a highly uniform volume flow with parallelly aligned streamlines (Fig. 1C) throughout the central part of the organoid cultivation compartment (SI Fig. 1B). At a level difference of 3 mm (equivalent to approx. 30 Pa pressure difference), an average flow velocity of 0.71 μm/s was observed, indicative of an interstitial fluid flow regime (Fig. 1C). Based on the established CFD model, an initial reservoir pressure difference of 3 mm was selected to drive interstitial fluid flow through the hydrogel and thus provide optimal culture conditions for the embedded organoid. Over a period of 24 hours, hydrostatic pressure-driven flow profiles followed an exponential decay resulting in a 1.8-fold higher influx during the first 6.5 hours of dynamic exposure (SI Fig. 1D). This behavior correlates well with recent findings that the glymphatic system is under circadian control, featuring significantly higher glymphatic influx and thus clearance during resting compared to active phases.^42,43^

### 2.2 Dynamic cultivation significantly reduces the necrotic core formation and improves neuronal differentiation

Initial on-chip studies, employing healthy midbrain hMOs focused on assessing the effects of a dynamic cultivation milieu on both nutrient supply as well as tissue differentiation. To that end, WT hMOs were introduced either into the microfluidic device (see Fig.2A) based on the workflow depicted in Figure 1D or cultivated in a conventional static tissue culture plate set-up. Neurite outgrowth is essential for the formation of mature neuronal networks and can therefore be considered an important indicator of tissue differentiation. As depicted in Figure 2D/F, exposure to interstitial fluid flow rates significantly enhanced neurite outgrowth, with a 1.5-fold higher average maximum neurite outgrowth rate in dynamically cultivated microtissues compared to static controls already 24 hours into the on-chip cultivation. Since insufficient nutrient supply in organoids is generally linked to the formation of so-called “dead cores,” WT hMOs were analyzed for the apoptotic marker caspase 3 after 35 days of differentiation. Strikingly, a 2-fold reduction in the normalized caspase 3 signal was observed in the presence of interstitial flow conditions (Fig. 2E/G), strongly pointing at improved nutrient availabilities under convective mass transfer.

**Figure 2:**
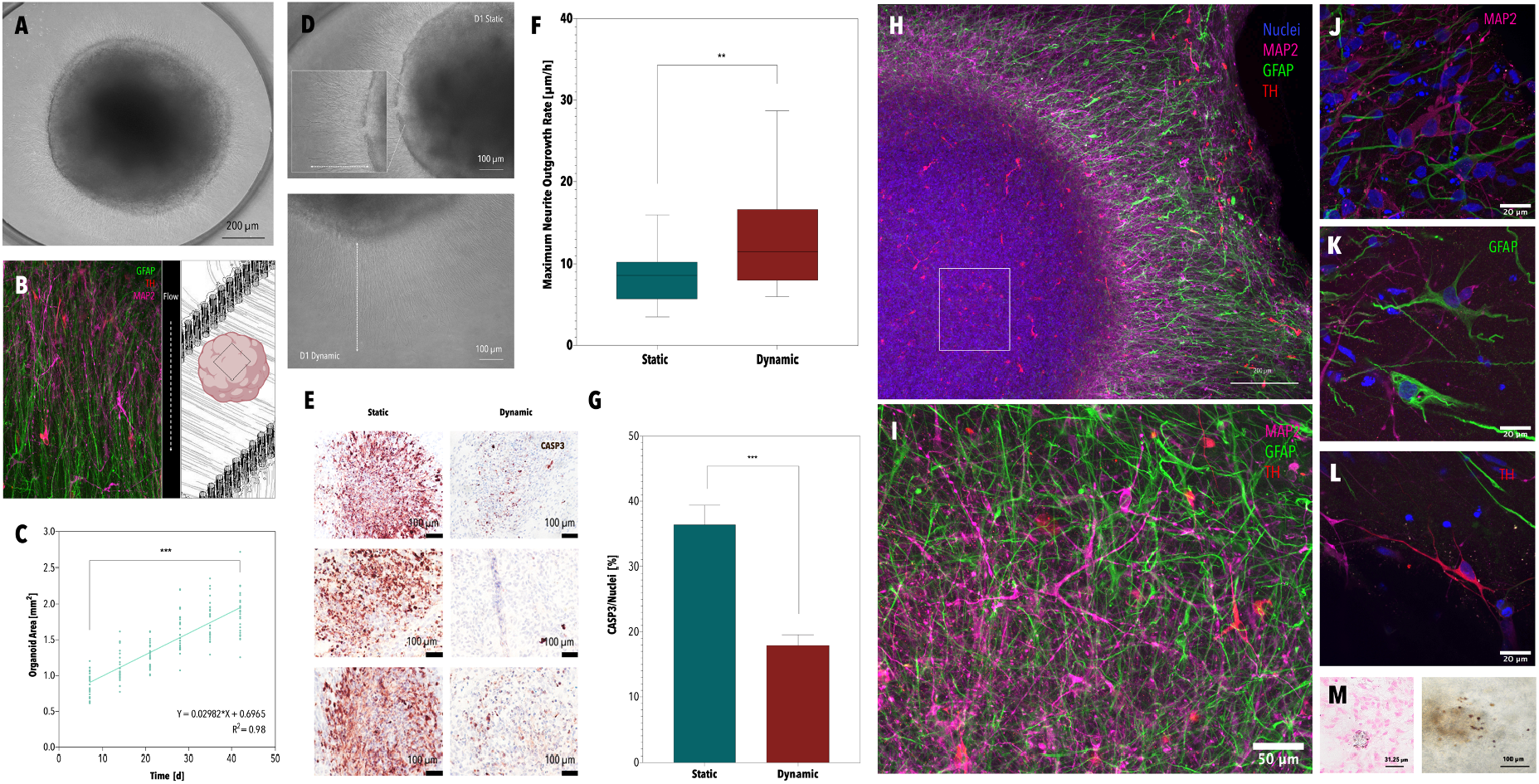
Brightfield image of an embedded hMO on-chip (A). Alignment of glial and neuronal processes in the direction of the applied flow: TH (red), GFAP (green), MAP2 (magenta) (B). Growth curve of midbrain microtissues on-chip. Statistical significance by mixed-effects analysis and Tukey test *p<0.033, **p<0.002, ***p<0.001 (n=8-10 from 3 independent organoid generations) (C). Brightfield images of statically (top panel) and dynamically (bottom panel) cultivated hMOs depicting differences in neurite outgrowth (left panel) (D). Boxplot of maximum neurite outgrowth rates of statically and dynamically cultivated hMOs. Statistical significance by Mann-Whitney test *p<0.033, **p<0.002, ***p<0.001. (n >= 3, from 3 independent organoid generations) (F). Micrographs and the respective quantitative analysis of immunohistochemically stained sections of hMOs depicting significant differences in the apoptotic marker caspase 3 after 35 days of differentiation. Statistical significance by Welch t-test *p<0.033, **p<0.002, ***p<0.001. Column and error bars represent mean±SEM (n >= 3, from 3 independent organoid generations) (E,G). Whole-mounted midbrain organoid after 60 days of differentiation: TH (red), GFAP (green), MAP2 (magenta), nuclei (blue) (H). Enlarged detail of the core of the whole-mounted hMO (H) at a magnification of 60x (I). Immunofluorescence staining of MAP2-positive neurons (J). Immunofluorescence staining of GFAP-positive astrocytes (K). Immunofluorescence staining of TH-positive dopaminergic neurons (L). Brightfield image (right panel) of neuromelanin aggregates in a midbrain organoid and the corresponding Fontana Masson staining revealing intra- and extracellular neuromelanin aggregation (left panel) (M).

Overall, morphological assessment of dynamically cultivated microtissues revealed a significant increase in hMO size over time, ranging from 0.87 ± 0.17 mm^2^ at day 7 to 1.87 ± 0.3 mm^2^ at day 42 (Fig. 2C). In addition, immunofluorescence analysis of WT hMOs after 50 days in dynamic culture revealed the presence of key midbrain-associated cell types including microtubule-associated protein 2 (MAP2)-positive neurons (Fig. 2J), tyrosine hydroxylase (TH)-positive dopaminergic neurons (Fig. 2L) as well as glial fibrillary acidic protein (GFAP)-positive astrocytes (Fig. 2K). Unexpectedly, a unidirectional interstitial fluid flow resulted in the alignment of both neuronal and glial cellular processes, clearly reflecting the parallel arrangement of the simulated streamlines as seen in Figures 1C and 2B. Occasionally, neuromelanin-like granules were detected within hMOs on-chip starting at about 30 days of differentiation (Fig. 2M/SI Fig.2A). Fontana-Masson staining confirmed the presence of both extra- and intra-cellular neuromelanin deposits, indicating that this physiological pigment, which is inherent to the *substantia nigra*, gets secreted within hMOs on-chip. To further support the observation that interstitial fluid flow improves midbrain tissue differentiation a comparative whole-mount analysis of WT hMOs was conducted after 50 days of on/off-chip culture. In addition to robust neuronal differentiation, dynamically cultivated microtissues displayed substantial glial differentiation, characterized by the presence of highly ramified GFAP-positive astrocytes. Moreover, immunofluorescence analysis of whole-mounted hMOs revealed a 2.1-fold higher TH/MAP2 ratio, as well as a 1.4-fold higher GFAP signal for dynamically cultivated hMOs compared to static controls (SI Table 1), pointing towards improved microtissue differentiation on-chip. In summary, the introduction of hMOs into our microfluidic platform not only supported tissue differentiation but also resulted in a marked improvement of one of the key limitations in organoid technology: insufficient nutrient supply.

### 2.3 Patient-specific hMOs (3×SNCA)-on-chip recapitulate key pathological hallmarks of PD

Following the establishment of a healthy microphysiological midbrain model, organoids derived from a PD patient carrying a triplication mutation of the α-synuclein gene were added to the study. A combination of morphological and immunofluorescence analysis was performed to verify the presence of PD phenotypes. Figure 3A shows the organoid growth dynamics of healthy and PD hMOs featuring distinct differences over a seven-week on-chip cultivation period. While WT (healthy) hMOs followed a linear trend that plateaued at around day 42, 3×SNCA hMOs displayed a complex dynamic growth behavior that can be characterized by an initial steep incline (D7-14), followed by a steady increase that subsequently declined by day 42. Whereas the organoid growth rates of healthy hMOs did not markedly change during the first 35 days of cultivation (0.033 ± 0.01 mm^2^/ day), the growth rates of PD hMOs displayed a progressive decline over time ranging from 0.082 mm^2^/ day on day 14 to 0.031 mm^2^/ day at day 35 (SI Fig. 3A). Upon comparison to their healthy controls, PD hMOs exhibited a 1.3-fold bigger cross-sectional area, with an up to 2.3-fold higher growth rate in the first phase of cultivation.

**Figure 3:**
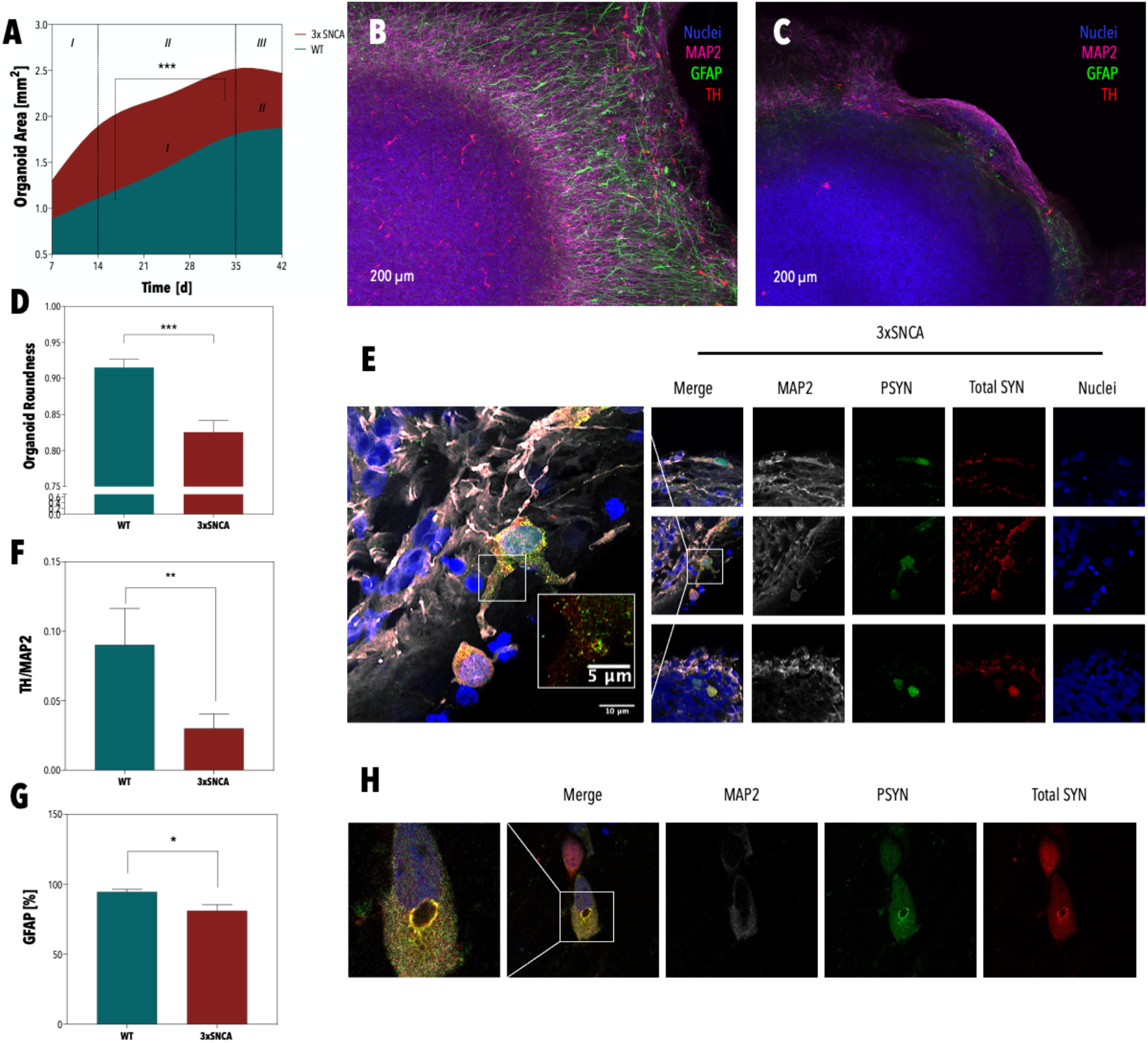
Growth curve of healthy and PD hMOs on-chip. Statistical significance by mixed-effect analysis and Tukey test *p<0.033, **p<0.002, ***p<0.001 (n=8-10 from 3 independent organoid generations) (A). Image of an immunofluorescence-stained healthy hMO after 60 days of differentiation on-chip: TH (red), GFAP (green), MAP2 (magenta), nuclei (blue) (B). Image of an immunofluorescence-stained PD hMO carrying a triplication mutation of the α-synuclein gene after 60 days of differentiation on-chip: TH (red), GFAP (green), MAP2 (magenta), nuclei (blue) (C). Comparative analysis of organoid roundness of WT and 3xSNCA hMOs at D31 of differentiation. Statistical significance by Welch’s t-test *p<0.033, **p<0.002, ***p<0.001. Column and error bars represent mean ± SEM (n=8-10 from 3 independent organoid generations) (D). Micrographs of 100 μm-thick sections of PD hMOs depicting the colocalization of α-synuclein and p-S129-a-synuclein after 60 days of differentiation on-chip (E). Quantitative analysis of immunofluorescence-stained WT and PD hMOs revealing significantly reduced levels of TH/MAP2 ratios. Statistical significance by Mann-Whitney test *p<0.033, **p<0.002, ***p<0.001. Column and error bars represent mean ± SEM (n=6-8 from 3 independent organoid generations) (F) Quantitative analysis of immunofluorescence-stained WT and PD hMOs revealing significantly reduced levels of GFAP-positive astrocytes. Statistical significance by Mann-Whitney test *p<0.033, **p<0.002, ***p<0.001. Column and error bars represent mean ± SEM (n=6-8 from 3 independent organoid generations) (G). Lewy-body-like inclusions observed in 100 μm-thick sections of 3xSNCA hMOs after 60 days of differentiation on chip (H).

The observation that 3×SNCA hMOs displayed a rapid size increase early on followed by a reduction in cell mass after 42 days correlates well with our previous findings that neurogenerative degeneration in 3×SNCA hMOs is preceded by an overproduction of the dopaminergic neuron population at early stages of differentiation.^44^ Figure 3D further shows that 3×SNCA hMOs displayed aspherical morphologies, as characterized by a significant reduction in organoid roundness from 0.92 for WT hMOs to 0.83 for 3×SNCA hMOs. Most importantly, immunofluorescence analysis of both healthy controls and 3×SNCA hMOs after 60 days of differentiation revealed a significant (3-fold) reduction in TH-positive dopaminergic neurons within patient-specific microtissues (Fig. 3F) – indicative of a neurodegenerative phenotype. Moreover, a significantly lower number of GFAP-positive astrocytes was detected in PD hMOs, in line with previously reported pathology-associated impairments in glial differentiation.^44^ Further analysis of PD hMOs (Fig. 3E) revealed the presence of both α-synuclein as well as p-S129-α-synuclein – a post-translationally modified version, which was found to be enriched in newly formed aggregates *in vivo*.^45^ Colocalization of p-S129-α-synuclein and α-synuclein was validated by calculating the Pearson’s correlation coefficient (>0). Intriguingly, immunofluorescence analysis additionally pointed towards the presence of Lewy body-like inclusions (the second hallmark of PD), as characterized by a peripheral halo, enriched in both α-synuclein and p-S129-α-synuclein and an electron-dense core (Fig. 3H and SI Fig. 3C).^46^ This means that two key pathological hallmarks of PD, namely neurodegeneration and α-synuclein aggregation in the form of so-called Lewy-bodies, were observed after 60 days of dynamic culture.

### 2.4 Orthogonal sensing enables the time-resolved monitoring of PD-related phenotypes

To demonstrate the analytical accessibility of our microfluidic midbrain organoid model, three non-invasive orthogonal sensing strategies (Fig. 1A) were evaluated in subsequent experiments to follow the onset and progression of (patho)physiological phenotypes. Figure 4A shows the position and working principle of the luminescence-based oxygen microsensors integrated into the microfluidic platform. Monitoring of healthy hMOs revealed a clear correlation between organoid growth and respiratory activity within the first 21 days of cultivation, resulting in an average normalized oxygen demand of 119.9 ± 4.4 hPa/mm^2^ (Fig. 4B). Comparative analysis of both healthy and 3×SNCA hMOs, shown in Figure 4D, revealed a significantly higher respiratory activity in PD microtissues over a cultivation period of 7 weeks. Similar to the complex growth behavior seen above (Fig. 3A), the oxygen demand of PD hMOs displayed high initiating demands of 137.7 ± 30.4 hPa, plateauing around 164.7±4.1 hPa at day 21, before eventually dropping to 155.1 ± 10.5 hPa around day 42. These results align well with our previous findings that neurodegenerative processes in 3xSNCA hMOs are preceded by an overproduction of the dopaminergic neuron population (e.g., high mitochondrial activity) early on.^44^ In turn, healthy hMOs displayed a steady increase in oxygen demand from 112.6 ± 35.85 hPa at day 3 to a peak of 159.7 ± 17.8 hPa at day 21, followed by a subsequent decline in respiratory activity between day 21 and day 42. This dynamic respiratory profile of healthy hMOs might reflect the chronology of brain development where neurogenesis (e.g., high mitochondrial activity) precedes astrogenesis (e.g., glycolytic metabolism).^47,48^ In an attempt to account for cell number variations between healthy and PD hMOs, organoid size-corrected data was calculated, revealing an inverted respiratory behavior. Results in Figure 4F point towards a significantly lower normalized oxygen demand in PD microtissues, indicative of impaired cellular respiration. To verify whether the reduction in normalized oxygen demand in PD hMOs can be linked to increased levels of aggregated α-synuclein, which has previously been shown to bind to and impair mitochondria,^49^ a comparative immunofluorescence analysis was performed. Quantitative image analysis using an in-house-developed algorithm confirmed elevated levels of p-S129-α-synuclein in 3×SCNA hMOs in relation to healthy controls (Fig. 4G). To subsequently ensure that reduced normalized oxygen demands of 3xSNCA hMOs are caused by impaired mitochondrial physiology, fluorescence analysis employing a TOM20 antibody was performed. Indeed, distinct differences encompassing a significant reduction in the overall mitochondria count, reduced mitochondrial complexity, as well as lower mitochondria numbers within dopaminergic neurons were detected for PD hMOs (Fig. 4E/C).

**Figure 4:**
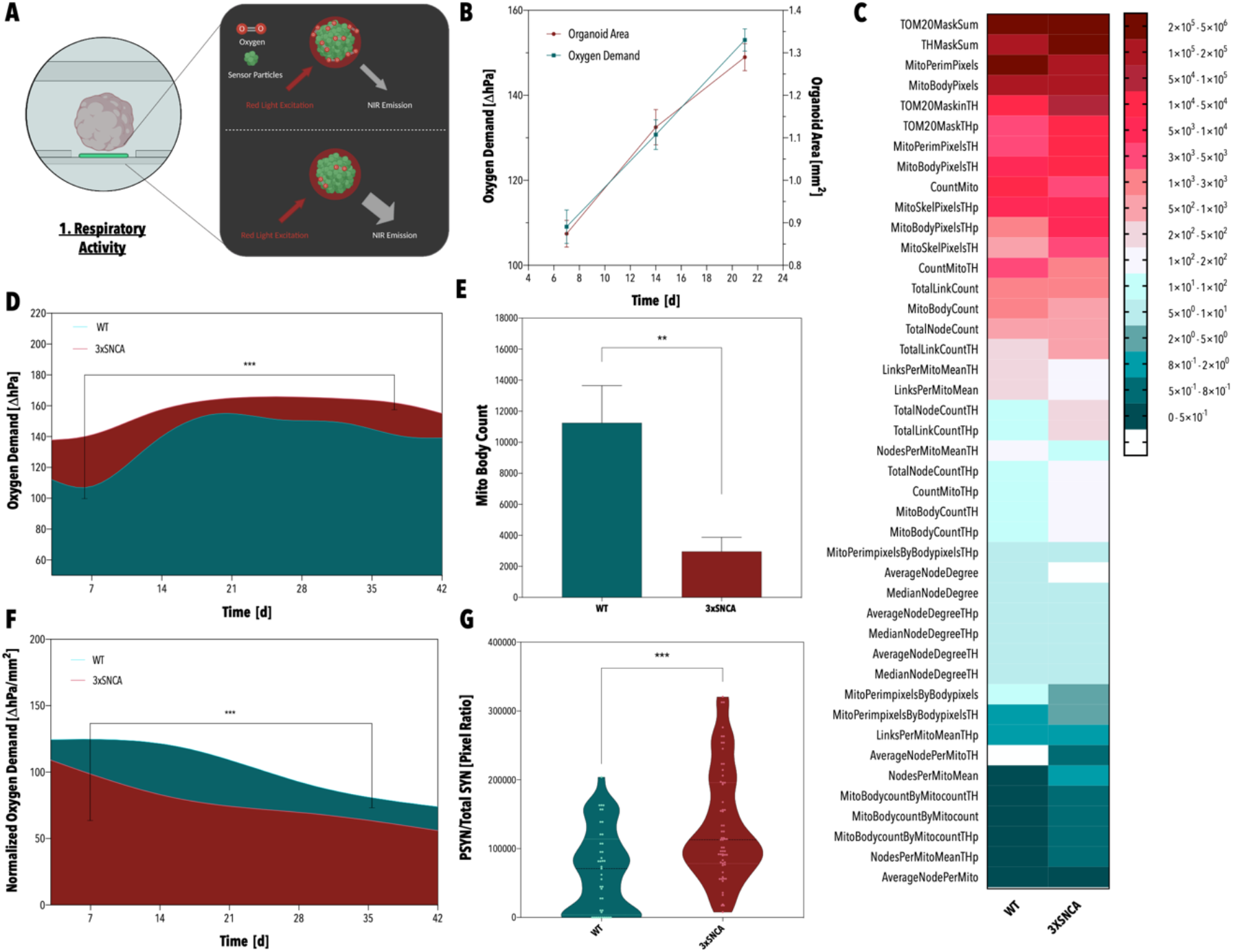
Schematic of the arrangement and working principle of the optical oxygen sensor (A). Correlation between healthy hMO growth and oxygen demand over 21 days of on-chip cultivation (B). Violin plot revealing elevated p-S129-α-synuclein to α-synuclein ratios in 3xSNCA hMOs (G). Heatmap providing an overview of altered mitochondria markers observed in 3xSNCA hMOs. (n >= 3 from 3 independent organoid generations) (C). Fitted oxygen demand profiles of healthy and PD hMOs over a cultivation period of 42 days (D). Fitted normalized oxygen demand profiles of healthy and PD hMOs over a cultivation period of 42 days (F). Comparative analysis of healthy and PD hMOs revealing significantly reduced mitochondria body counts in 3xSNCA hMOs. Column and error bars represent mean ± SEM (E). (D,F) Statistical significance by mixed-effect analysis and Tukey test *p<0.033, **p<0.002, ***p<0.001 (n=8-10 from 3 independent organoid generations). (E,G) Statistical significance by Mann-Whitney test *p<0.033, **p<0.002, ***p<0.001 (n >= 3 (G) n >= 3 from 3 independent organoid generations (E)).

The neurotransmitter dopamine can be considered a crucial PD-specific biomarker, as its levels directly reflect the tissue’s pathological status. By inserting a nanoparticle-based enzymatic biosensor into the waste reservoir of the microfluidic device, changes in dopamine concentrations in the organoid’s supernatant were recorded over time. Figure 5A provides a schematic of the hydrogel-covered carbon electrode, comprising the enzyme tyrosinase, the hydrogel chitosan, and the two oxygen donors, cerium oxide and titanium oxide. Results in Figure 5B show a steady increase in dopamine levels for both healthy and 3×SNCA hMOs over a period of 7 weeks, with the first detectable signals retrieved after 35 days of microfluidic cultivation. By week 7 (59 days of differentiation), significant differences in dopamine secretion between healthy and PD hMOs started to emerge, correlating well with our previous findings that showed first discernable variations in TH expression around day 60 of differentiation.^44^ In addition, the electrochemical analysis supported our earlier observation that dynamic cultivation improves tissue differentiation, or the TH-positive neuronal population, respectively, with a 3.9-fold higher dopamine signal obtained for hMOs exposed to interstitial fluid flows compared to static controls (Fig. 5C). Since electrophysiological activity constitutes an important (patho)physiological parameter, assessing neuronal firing activity is essential for any *in vitro* brain model. Consequently, a multielectrode array (MEA) was integrated into the supply and waste channels of the organ-on-a-chip platform, enabling electrophysiological recordings of the neuronal microtissues (Fig. 5H/D/E). Flow-directed outgrowth was utilized to direct neuronal processes extending from the three-dimensional microtissue onto the two-dimensional electrode array. (Fig. 5E/H). By using this approach, spontaneous electrophysiological activity was recorded starting at 24 days of differentiation (Fig. 5G). Next to monophasic and biphasic spikes (Fig. 5Ib/c), various firing patterns were observed, including tonic spiking, phasic bursting as well as tonic bursting (Fig. 5F). Overall, 66 ± 14 % of all active electrodes displayed bursting activity, a characteristic of dopaminergic neurons.^50^ Dopaminergic identity was further supported by clusters of spikes that displayed breaks in the initial spike segments (Fig. 5Ia).^51^ Electrophysiological activity was validated by silencing with the neurotoxin tetrodotoxin (Fig. 5F) as well as by performing Fluo-4 acetoxymethyl ester (AM)-based calcium imaging (data not shown).

**Figure 5:**
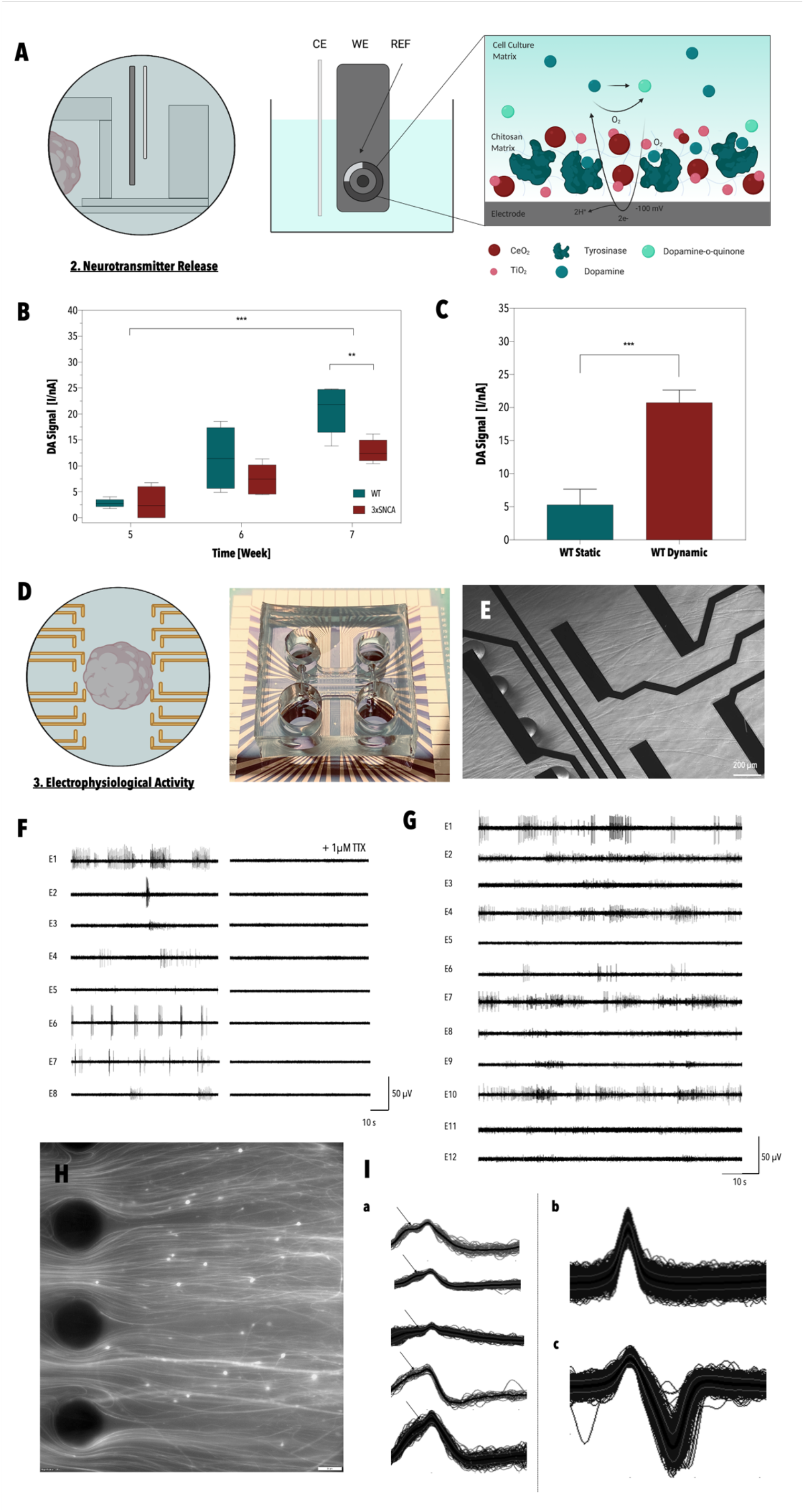
Schematic of the working principle of the electrochemical dopamine biosensing approach and the three-electrode set-up (A). Increase in dopamine signal in the supernatant of healthy and PD midbrain organoids over time. Statistical significance by two-way ANOVA and Tukey test *p<0.033, **p<0.002, ***p<0.001 (n=10 (pooled) from 2 independent organoid generations (technical triplicates)). Comparative analysis between WT and 3xSNCA signals (week 7) was conducted by Welch ‘ t-test *p<0.033, **p<0.002, ***p<0.001 (n=10 (pooled) from 2 independent organoid generations) (B). Comparative analysis of the dopamine signal of statically and dynamically cultivated WT hMOs. Statistical significance by Welch ‘ t-test *p<0.033, * *p<0.002, * * *p<0.001 Column and error bars represent mean ± SEM (n=10 (pooled) from 2 independent organoid generations (technical triplicates)) ((C). Schematic and image of the MEA integrated microfluidic device (D). Micrograph of neurites extending over the microelectrodes on-chip (E). Electrophysiological recordings of human midbrain organoids on-chip prior to and after the exposure to the neurotoxin TTX (F). Representative read-out of the electrophysiological activity of a healthy hMO in a microfluidic device (G). Flow-directed outgrowth of neurites on-chip (H). Examples of dopaminergic clusters, characterized by a break in the initial segment (Ia/arrow) as well as mono- and bi-phasic spikes recorded on-chip (I_b/c_).

### 2.7. On-chip monitoring reveals HP-β-CD-mediated phenotypic rescue in 3xSNCA organoids

While HP-β-CD has been established as a promising excipient in pharmacology, its repurposed application for enhancing autophagic capacity in PD organoids has only been reported recently.^39^ In an attempt to demonstrate the ability of our multi-sensor integrated midbrain organoid-on-a-chip platform to detect phenotypic rescue, exposure of 3×SNCA hMOs to 5 μM of HP-β-CD was assessed. To that end, dopamine release and respiratory activity of healthy, 3×SNCA, as well as HP-β-CD-treated 3xSNCA hMOs were investigated in a final time-resolved comparative study. As autophagy is directly involved in the removal of both aggregated protein species as well as impaired organelles, specific focus has been directed towards assessing mitochondrial parameters and p-S129-α-synuclein to α-synuclein ratios. While no significant differences in dopamine release were found between 3xSNCA hMOs and HP-β-CD treated 3×SNCA hMOs (Fig. 6E), markedly altered growth behaviors were observed. Figure 6G shows a significantly lower mean organoid growth rate of 0.02 μm^2^/day for treated hMOs compared to 0.03 μm^2^/day for untreated controls (D14-D35), resulting in overall smaller microtissues upon treatment with HP-β-CD. As improved autophagy has been linked to limited cell growth, we speculate that the average size reduction of 12% in HP-β-CD treated hMOs after 42 days of on-chip cultivation is indicative of enhanced cellular degradation.^52^ Furthermore, when taking into account reduced hMO sizes into the DA measurements, by normalizing the amperometric signal to microtissue size, CD treated hMOs give rise to an average 1.73-fold higher DA signal compared to their untreated controls. While the highest overall oxygen demand of 102.8 ± 31 hPa/mm^2^ (n=410) was detected in healthy hMOs, HP-β-CD treatment significantly raised the average normalized oxygen demand of PD hMOs from 78.21 ± 23.2 hPa/mm^2^ (n=587) to 88.54 ± 21.8 hPa/mm^2^ (n=541), pointing at improved mitochondrial physiology. To confirm that the observed phenotypic improvements originate from rescue effects of PD-associated phenotypes, such as α-synuclein aggregation, additional immunofluorescence analysis was performed after 60 days of differentiation. As shown in Figure 6C, quantitative analysis of hMO-sections revealed a significant reduction in the amount of pathological p-S129-α-synuclein, with comparable p-S129-α-synuclein to α-synuclein ratios in healthy and HP-β-CD treated hMOs. These observations support previous findings that impaired autophagic clearance promotes the exocytosis of α-synuclein, ultimately linking autophagic dysfunction to a progressive spread of Lewy pathology.^53^ To assess whether reduced levels of aggregated p-S129-α-synuclein consequently resulted in improved mitochondrial morphology, fluorescence analysis was performed. In agreement with the p-S129-α-synuclein data, we observed marked rescue effects in mitochondrial physiology for HP-β-CD treated hMOs, as indicated by the clustering of the WT and the CD-treated group in a hierarchical cluster analysis (Fig. 6D). In a final attempt to prove, the beneficial effects of HP-β-CD, a whole-mount analysis (Fig. 6H) was performed. Image analysis revealed a significant increase in dopaminergic neurons (TH/MAP2 ratio), the afflicted cell type in PD, following treatment with the repurposed excipient HP-β-CD. This finding supports our previous assumption that elevated DA levels, were masked by reduced microtissue sizes observed in HP-β-CD treated organoids.

**Figure 6:**
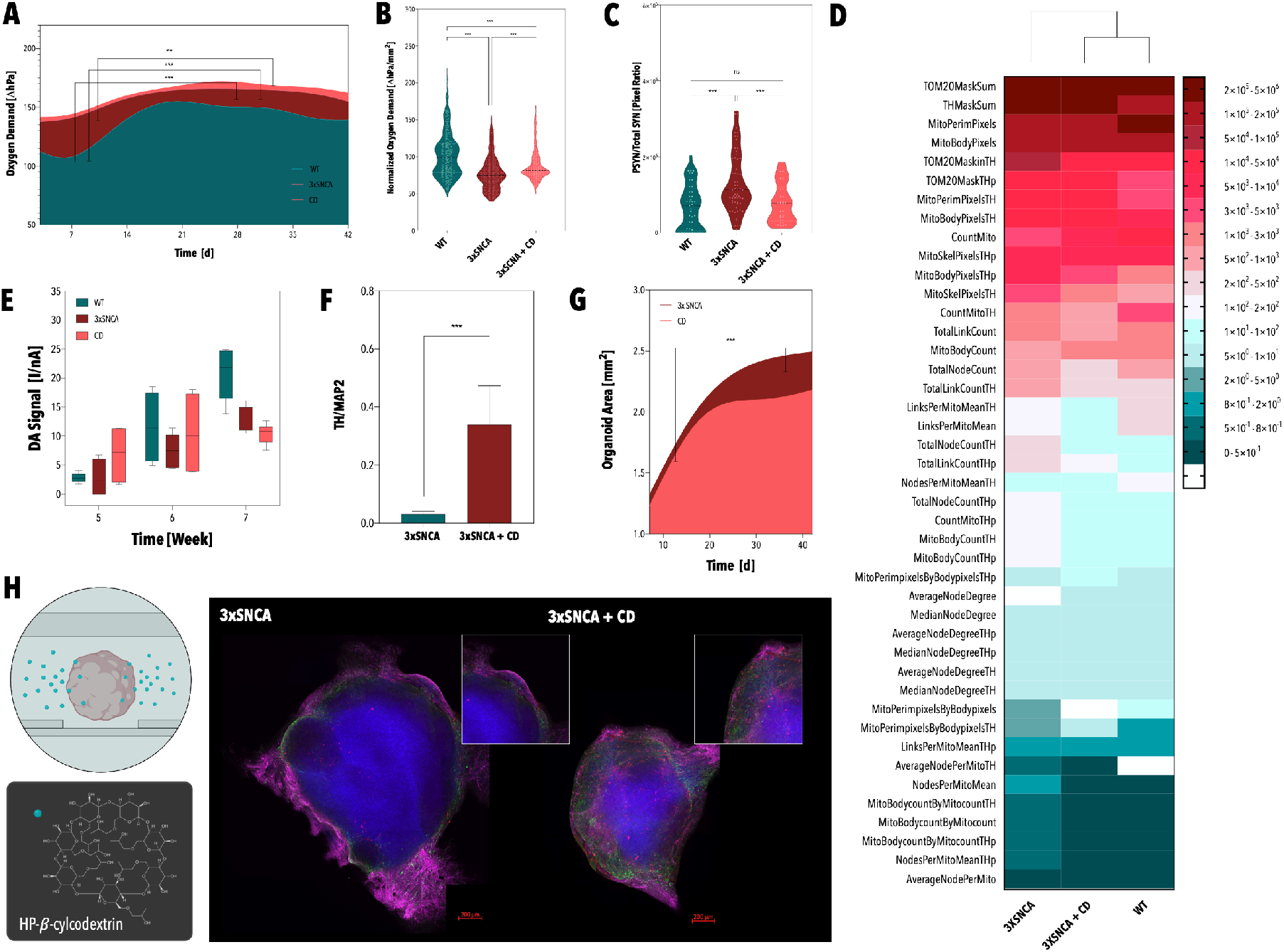
Oxygen demand profiles of healthy, 3×SNCA, and HP-β-CD-treated hMOs over a cultivation period of 42 days (A). Normalized oxygen demands of healthy, 3×SNCA, and HP-β-CD-treated hMOs. Statistical significance by Kruskal-Wallis test and Dunn’s multiple comparison test *p<0.033, **p<0.002, ***p<0.001 (n=8-10, from 3 independent organoid generations and 12 timepoints) (B). Violin plot of p-S129-α-synuclein to α-synuclein ratios calculated for healthy, 3xSNCA, and HP-β-CD-treated hMOs. Statistical significance by Kruskal-Wallis test and Dunn’s multiple comparison test *p<0.033, **p<0.002, ***p<0.001 (n >= 3) (C). Heatmap-based overview of improved mitochondrial markers in HP-β-CD-treated 3×SNCA hMOs. Hierarchical clustering was performed using MATLAB. (D) Time-resolved dopamine data of healthy, 3xSNCA, as well as HP-β-CD-treated hMOs, cultivated in microfluidic devices. Statistical significance by two-way ANOVA and Tukey test *p<0.033, **p<0.002, ***p<0.001 (n=10 (pooled) from 2 independent organoid generations (technical triplicates) (E). Comparative image analysis of immunofluorescence-stained 3×SNCA and HP-β-CD-treated 3xSNCA organoids revealing a significant rescue in the TH/MAP2 ratio. Statistical significance by Mann-Whitney test *p<0.033, **p<0.002, ***p<0.001. Column and error bars represent mean ± SEM (n> = 5 from 3 independent organoid generations) (F). Growth curves of 3xSNCA hMOs and HP-β-CD-treated 3xSNCA hMOs over a cultivation period of 42 days (n=8-10 from 3 independent organoid generations) (G). Graphical illustration of HP-β-CD-treatment and representative fluorescence images of a 3×SNCA hMO (left panel) and a HP-β-CD-treated 3xSNCA hMO (right panel): TH (red), GFAP (green), MAP2 (magenta), nuclei (blue) (H). (A,G) Statistical significance by mixed-effect analysis and Tukey test *p<0.033, **p<0.002, ***p<0.001 (n=8-10 from 3 independent organoid generations).

## 3. Discussion and Conclusion

In this study, we have developed a patient-specific multi-sensor integrated midbrain organoid-on-a-chip model capable of monitoring the onset, progression, and rescue of PD-related phenotypes, while simultaneously providing for shorter organoid handling times, reduced contamination risks as well as high content analysis compatibility.^54, 8^ It is important to note that the quality of any *in vitro* disease/tissue model can be defined by two parameters 1) the (patho)physiological similarity to the organ of interest and 2) its accessibility to biochemical analysis methods. Both aspects are essential to foster clinical translation in precision medicine and routine application in pharmaceutical development. In this work, we have taken advantage of the synergistic effects created by combining organoid technology and its capability to emulate intricate organotypic structures *in vitro* with organ-on-a-chip systems that allocate analytical accessibility, scalability, and physiological fluid dynamics. A key biophysical aspect of our multi-sensor integrated midbrain organoid-on-a-chip platform is the application of flow profiles that mimic brain-specific circadian flow regimes, resulting in a drastic reduction of necrotic core formation - a key limitation of organoid technology - and improved midbrain tissue differentiation.^55^ In other words, cultivating hMOs under interstitial fluid flow clearly promotes brain organogenesis by increasing oxygen availabilities, improving nutrient supply, and promoting the removal of locally accumulated toxic compounds such as acids or reactive oxygen species.^56^ Analytical accessibility through non-invasive multi-parametric sensing strategies is particularly important when studying time-dependent processes, such as tissue differentiation or disease progression, in an otherwise contained *in vitro* system. Time-resolved oxygen sensing of healthy hMOs, for example, reflected the sequential nature of brain development, characterized by high initial neuronal differentiation (correlating with increasing respiratory activities) that is succeeded by astrogenesis (marked by lower respiratory activities) initiated at about day 21 of hMO differentiation.^44,57^ In accordance with the degree of tissue differentiation, electrochemical dopamine measurements revealed a time-dependent increase in dopamine signals with significantly higher catecholamine levels observed under dynamic cultivation conditions compared to static controls. Electrophysiological recordings further supported functional network maturity and the presence of dopaminergic populations. A comparative analysis between healthy and PD hMOs (3×SNCA) within our multi-sensor integrated microfluidic platform revealed a clear time dependency in the onset of Parkinson’s disease-related phenotypes, reflecting the complex progression of the neurodegenerative disorder. While initial high numbers in TH-positive neurons can be attributed to differentiation upregulating effects elicited by elevated levels of α-synuclein,^33^ heightened dopamine metabolism, in turn, was shown to promote oxidative stress that together with the accumulation of toxic forms of α-synuclein subsequently can contribute to increased neurotoxicity and -degeneration.^58^ Importantly, elevated levels of p-S129-α-synuclein, impaired mitochondrial physiology, diminished TH-positive neuronal populations, as well as the presence of Lewy-body-like inclusions, were detected in 3xSNCA organoids, demonstrating the validity of our microfluidic *in vitro* model for studying PD.^46^ Since the observed pathological phenotypes can be linked, at least in part, to impaired autophagy, the repurposed excipient HP-β-CD, which has previously been shown to induce rescue effects in a PD organoid model with mutations in PINK1, was used to challenge the ability of our multi-sensor integrated midbrain organoid-on-a-chip platform.^39^ Intriguingly, integrated oxygen sensing alone pointed towards significant rescue effects expressed by a marked increase in normalized oxygen demands for HP-β-CD-treated hMOs compared to PD controls. These effects were further verified by immunofluorescence analysis, revealing significant reductions in aggregated α-synuclein, distinct improvements of several mitochondrial characteristics as well as significantly enlarged populations of TH-positive dopaminergic neurons.

Overall, the presented multi-sensor integrated platform significantly improved the differentiation of midbrain microtissues by exposing them to physiological flow profiles and enabled for the first time in an on-chip brain organoid model the ability of non-invasive and multi-parametric monitoring. As such, it surpasses existing in-vitro models of the human midbrain, opening the way to precision medicine employing personalized PD models.

## Materials and Methods

### Microfabrication

The microfluidic chip was designed using the CAD program Autocad. Pre-molds were fabricated by casting polydimethylsiloxane (PDMS/ 1:10 ratio) (Sylgard®184 Silicone Elastomer Kit, Down Corning) from a computer numerical control (CNC) milled polycarbonate master mold using standard soft lithography techniques (SI Fig. 1C). After 2 h of polymerization at 80 °C, the pre-mold was removed from the polycarbonate master mold and cured for another 24 h at 120°C. Afterward, the pre-mold was silanized with 2 μL of trichloro (1H, 1H, 2H,2H-perfluorooctyl) silane (Sigma Aldrich) and incubated for 1 h at 80°C. The same procedure was repeated for the generation of the PDMS mold, starting from the silanized pre-mold (SI Fig. 1C). PDMS molds were used for the fabrication of the microstructured top layers. To facilitate the removal of the microstructured layer from the mold, ethanol (absolute, ChemPur) was applied between the two layers prior. After 2h of polymerization at 80 °C, microfluidic reservoirs and inlets were introduced employing biopsy punches (Ø 6mm, Ø 8mm, Ø 2mm, Kai Medical). Cleaned microstructured PDMS top layers (adhesive tape (Scotch)) were subsequently bonded to blotted 250 μm PDMS foils as well as cleaned glass substrates using air plasma (Harrick Plasma, High Power, 2 min). PDMS foils (MVQ Silicones GmbH) were processed using xurography (CAMM-1 GS-24, Roland). Microfluidic devices equipped with oxygen sensors were generated by the deposition of 2 μL of a microparticle solution into PDMS cavities within glass substrates by the use of a pipette as previously described by *Zirath et al*.^30,31^ After drying for 2 h at room temperature, the microparticles were immobilized to the glass substrate, and the fluidic structures were sealed, employing air plasma. Prior to their use, microfluidic devices were sterilized, utilizing a combination of 70 % ethanol as well as UV treatment.

### CFD Simulation

A multipurpose finite volume CFD code (Ansys Fluent 6.3.26, www.ansys.com / OpenFoam www.openfoam.org) was used for calculating the flow profile on-chip. The geometry consisting of the hydrogel cavity, the two feed channels, and the two collection units was split into 136.000 hexahedral control volumes (SI Fig. 1A). The grid pillars at the gel inflow and outflow boundary were fully resolved. For adequate numerical accuracy, second or higher-order discretization schemes have been selected for all flow variables (Navier-Stokes equation – momentum conservation, Continuity equation – mass conservation) and the species equations. All wall boundaries were treated as ideally smooth; no-slip boundary conditions (zero flow velocity at the wall) were selected for all surfaces. The outlet was set to a pressure outlet at a standard pressure of p = 1 atm (101325 Pa). The hydrogel region was approximated as a homogeneous and isotropic porous zone (Darcy-Forchheimer equation) with a constant porosity of ε = 0.99 and viscous resistance of R = 1.89·10^13^ 1/m^2^ having been assumed for all directions.^59–64^ Isothermal flow was assumed, no temperature or energy field was solved. For simplicity, Newtonian fluid behavior was applied for the simulation using a constant dynamic viscosity and constant density (incompressible) for all mixture components. As the concentrations of the dissolved species in the fluid are low, the properties of the solvent, water, have been used for the simulation (ρ = 993 kg/m^3^, η = 0.001003 Pa·s at 37 °C). The diffusion coefficients for the tracer components have been estimated according to literature values (glucose: 0.18 kDa – 4·10^-10^ m^2^/s, oxygen: 32 Da – 2·10^-9^ m^2^/s, water: 18 Da – 2·10^-9^ m^2^/s) assuming a dilute solution.^60^ Different water species have been used for both inlets to investigate the cross mixing of the two inlet channel fluids. Simulations were carried out on the cluster server cae.zserv.tuwien.ac.at (operated by the IT department of TU Wien, www.zid.tuwien.ac.at). As the major flow resistances are inside the hydrogel and in the flow channels, but not in the feed and collection cavities, a simplification was used: To reduce the computational effort, steady-state simulations for different selected feed cavity filling levels have been carried out. The simulated filling level was translated into a corresponding relative pressure difference between the feed inlet zone and the pressure outlet.

### hMO Generation and On-Chip Cultivation Protocol

iPSCs from a healthy individual (identifier: 2.0.0.51.0.0) and a PD patient (triplication mutation of 3xSNCA, identifier: 2.1.3.138.0.0) were used in this study. The maintenance of iPSCs was performed as previously described.^65^ From the iPSC line, human ventralized neural epithelial stem cells (hvNESC) were generated, which were subsequently used to generate midbrain organoids.^12^ For on-chip hMO cultivation, organoids suspended in Matrigel^®^ (Corning) were transferred into the microfluidic chip on day 0 of the maturation phase and cultivated for up to 60 days of differentiation. Dynamic cultivation was achieved by filling the feed medium reservoirs up to a 3.4 mm feeding level, while the medium at the collector side was kept at 0.4 mm height. The medium was exchanged every 48 - 72 h. To prevent any drying out of the microfluidic chips, (i) devices were kept in a compartmentalized cultivation platform (Quadriperm^®^, Greiner) with 10 mL of PBS on both sides, and (ii) inlets of the hydrogel chamber were sealed using PCR tape (Sigma Aldrich). Depending on the type of analysis, static controls were either cultivated in microfluidic devices under static conditions or were embedded in a droplet of Matrigel^®^ (Corning^®^) and cultivated in a 24-well plate (Greiner). Equal amounts of cell culture medium and Matrigel^®^ (Corning^®^) were used, and the medium change procedure was kept identical under static conditions.

### HP-β-CD-Treatment

For HP-β-CD-treatment 3×SNCA hMOs were exposed to 5 μM HP-β-CD (Sigma Aldrich) in cell culture media starting from D10 of differentiation.

### Immunohistochemistry

#### Chromogenic Immunohistochemistry

##### Caspase 3 Staining

HMOs were fixed with 4 % paraformaldehyde overnight at room temperature, washed three times with phosphate-buffered saline (PBS) (Gibco) for 15 minutes, before being dehydrated, embedded in paraffin, and sectioned using a microtome (Thermo Scientific HM355 S). Prior to heat-induced antigen retrieval using Tris-EDTA buffer at pH 9, sectioned hMOs were deparaffinized and rehydrated. After rinsing the sections in TBS, endogen peroxidase and alkaline phosphatase activity were blocked by treating the sectioned hMOs with BLOXALL (VectorLaboratories) for 10 min. The primary antibody (ASP175, #9661, CellSignaling) was incubated for one hour at room temperature before the sections were rinsed again in TBS and the secondary antibody, an anti-rabbit HRP conjugated antibody (BrightVision), was applied for 30 min at room temperature. Color development was achieved by a 6-minute-long exposure to ImmPACT™Nova Red™(VectorLaboratories). Subsequently, the sections were stained with Haematoxylin (Roth) and mounted with Epredia™ Consul-Mount™ (Fisher Scientific).

##### Fontana Masson Staining

HMOs were fixed with 4 % paraformaldehyde overnight at room temperature, washed three times with phosphate-buffered saline (PBS) (Gibco) for 15 minutes, before being dehydrated and embedded in paraffin, and sectioned using a microtome (Thermo Scientific HM355 S). After deparaffinization and rehydration, sectioned hMOs were stained for 10 minutes in a Lugol’s solution (2 g Potassium iodide (Roth), 1g Iodine (Roth)) before being transferred into a 5% sodium thiosulfate solution (Morphisto) for two minutes. Rinsed slides (3x, aqua dest.) were transferred in an ammoniacal silver solution (5% silver nitrate (Roth); ammonium hydroxide, 18% NH3 (Alfa Aesar)) and incubated at 60°C for two hours. After rinsing (3x, aqua dest.), slides were exposed to a 0,2% gold chloride solution (Fluka) for three minutes. Slides were rinsed in aqua dest. before being treated with 5% sodium thiosulfate solution for two minutes and subsequently rinsed in tap water for two minutes. Cell nuclei were stained using 0,1% nuclear fast red (Merck). After finishing the staining procedure, the slides were dehydrated in ethanol and mounted with Epredia™ Consul-Mount™ (Fisher Scientific).

#### Immunofluorescence

##### Sectioned hMOs

HMOs were fixed with 4 % paraformaldehyde overnight at room temperature and washed 3× with PBS for 15 min. For sectioned analysis, hMOs were embedded in 3-4 % low-melting-point agarose in PBS. The solid agarose block was sectioned with a vibratome (Leica VT1000s) into 100 μm sections. The sections were blocked for 90 min at room temperature on a shaker using 0.5 % Triton X-100, 0.1% sodium azide, 0.1% sodium citrate, 5 % normal goat serum, and 2% bovine serum albumin in trisbuffered saline (TBS). Primary antibodies were diluted in the same solution and incubated for 48 h at 4 °C. Antibodies were diluted according to the supplementary table SI Table 2. After incubation with the primary antibodies, sections were washed 3x with TBS for 15 minutes and incubated for 30 minutes in TBS supplemented with 0.5 % Triton X-100, 0.1% sodium azide, 0.1% sodium citrate, 5 % normal goat serum, and 2% bovine serum albumin. Subsequently, sections were incubated with the respective secondary antibodies (SI Table 2) and the nuclear dye Hoechst 33342 (Invitrogen) in TBS with 0.5 % Triton X-100, 0.1% sodium azide, 0.1%sodium citrate, 5 % normal goat serum, and 2% bovine serum albumin for 2 h at room temperature. Afterward, samples were washed 3x with TBS, 1× with Milli-Q water before being mounted in Fluoromount-G mounting medium (Southern Biotech). Sections were imaged using confocal microscopes (Yokogawa, Zeiss).

##### Whole-mounted hMOs

Paraformaldehyde fixed (4%) hMOs were washed 3x with PBS for 15 minutes on a shaker. Subsequently, hMOs were blocked and permeabilized using 1% Triton X-100 and 10% normal goat serum in 1× PBS for 24h on a shaker at room temperature. Primary antibodies were diluted in PBS supplemented with 0.5 % Triton X-100 and 3% normal goat serum and incubated for 4 days at 4°C on a shaker (SI Table 2). Samples were washed 3× with PBS for 1 hour at room temperature before incubating with the diluted secondary antibody solution at 4°C, including the nuclear dye Hoechst 33342 (Invitrogen). After 2 days, samples were washed 3× with 0.05% Tween-20 in PBS and 1× with Milli-Q water for 5 minutes at room temperature, prior to mounting with Fluoromount-G mounting medium (Southern Biotech). Images were acquired using a high content analysis system (Operetta^®^, PerkinElmer).

#### Oxygen Monitoring

On-chip oxygen monitoring was carried out at a sampling frequency of 1 Hz using a FireStingO_2_ optical oxygen meter (Pyroscience) connected to optical fibers (length 1 m, outer diameter 2.2 mm, fiber diameter 1 mm). Integrated sensors were calibrated using a CO_2_/O_2_ oxygen controller (CO_2_-O_2_-Controller 2000, Pecon GmbH) equipped with integrated zirconium oxide oxygen sensors (SI Fig. 4A/B). Oxygen measurements were performed every 3 to 4 days. For this purpose, chips were sealed with PCR foil and transferred into an external incubation chamber set-up (5% CO_2_ / 37°C). Each sample was measured for a minimum of 3 minutes to guarantee proper equilibration. To ensure that fluid flow does not interfere with the oxygen measurement, oxygen demand was measured prior to the reestablishment of flow (under static conditions). Oxygen demand was subsequently calculated according to the following formula: hMO oxygen demand (ΔP_O2_) = P_O2_ blank – P_O2_ hMO.

#### Dopamine Sensing

The fabrication and subsequent characterization of the dopamine sensor were based on a protocol previously published by Niagi *et al*.^66^ Carbon electrodes of a thick-film electrode set-up with an integrated silver reference electrode (S1PE, MicruX Technologies) were coated with 15 μL of a tyrosinase (Sigma Aldrich), chitosan (1%, Sigma Aldrich), CeO_2_ (10 mg/mL, Sigma Aldrich) and TiO_2_ (10 mg/mL, chemPUR) mixture at the following ratio (4:4:1:1) and incubated for a duration of 1 hour. Amperometric measurements were conducted at a potential of −0.15 V using a three-electrode set-up with a platinum wire as a counter electrode and a potentiostat (VMP3, Bio-Logic) equipped with a low current module (Bio-Logic). Dopamine (Sigma Aldrich) calibration curves were recorded in both PBS and N2B27 medium (SI Figure 4C/D). Concentrations of tested interferents were based on previous studies, media composition, and HPLC-derived neurotransmitter profiles of hMOs^67^. Next to 5 μM dopamine (dopamine hydrochloride), interference studies encompassed the measurement of 5 μM L-DOPA (Sigma Aldrich), 5 μM DOPAC (Sigma Aldrich), 5 μM norepinephrine (Sigma Aldrich), 5 μM epinephrine (Sigma Aldrich), 5 μM serotonin (Sigma Aldrich), 40 μM γ-aminobutyric acid (GABA) (Sigma Aldrich) and 200 μM ascorbic acid (Sigma Aldrich) in PBS (SI Figure 4F). To assess potential interference effects of phenol red on the DA measurement, a comparative analysis between phenol red basal media (Neurobasal Medium, Gibco) and phenol red-free basal media (Neurobasal Medium, minus phenol red, Gibco) employing 5 μM DA was conducted (SI Figure 4G). Stability measurements were performed at a DA concentration of 5 μM DA in PBS (SI Figure 4E). Measurements were conducted by dipping the three-electrode set-up into the respective analyte. For each measurement, the supernatants of 10 hMOs were pooled and three technical triplicates were recorded.

#### MEA Fabrication and Analysis of Electrophysiological Activity

The fabrication of the MEAs followed a previously published protocol by Mika *et al*.^68^ In detail, cleaned (sonication in acetone for 1 minute) glass substrates (49mm × 49mm glass substrates (D263 T ECO, Schott GmbH)) were sputtered with chrome using an RF power of 100W, 30s sputter time, a working pressure of 2×10^-5^ mbar and a base pressure of 8×10^-3^ mbar (Von Ardenne LS320 Sputtersystem). Photoresist AZ5214E was spun onto the chrome-coated substrates using a spin speed of 3000 rpm and a spin time of 30 s. The soft bake was performed at 100°C for 60 s. Using a Karl Suss MJB3 UV400 mask aligner, the mask and substrate were aligned and exposed to a dose of 40 mJ/cm^2^. Next, the reversal bake was performed at 120°C for 70 s and immediately flood-exposed to a dose of 240 mJ/cm^2^. After flood exposure, substrates were developed using AZ726 MIF for 60 s. Development was stopped by rinsing with water. After the substrates were dried with N2, the chrome layer was etched using CHROME ETCH 18 (Micro Resist Technology). After inhibiting the etching process by rinsing with water, substrates were dried. Titanium was sputtered with an RF power of 100 W, a 50 s sputter time, a working pressure of 2×10^-5^ mbar, and a base pressure of 8×10^-3^ mbar. Next, two layers of gold were sputtered with an RF power of 50 W, a sputtering time of 50 s, a working pressure of 2-5 mbar, and a base pressure of 8-3 mbar. Lift-off was performed by sonication in acetone, before rinsing the substrates with acetone and isopropanol. The remaining chrome was removed using CHROME ETCH 18 (Micro Resist Technology). The insulation layer of Si3N4 was deposited using plasma-enhanced chemical vapor deposition with an RF power of 12 W, a processing time of 30 minutes, 1 torr working pressure, 0.06 torr base pressure, a SiH4 flow of 700 sccm, an NH3 flow of 18 sccm and a substrate holder temperature of 300°C (Oxford Plasmalab 80 Plus). To etch the insulation layer, a sacrificial layer of AZ5214E was spun onto the substrates using a spin speed of 3000 rpm, a spin time of 30 s, and ramp set to 4. The soft bake was performed at 100°C for 60 s. The exposure dose was 40 mJ/cm^2^ (equals ~4s in the case of MJB3 Mask Aligner). Next, the reversal bake was performed at 120°C for 70 s and immediately flood-exposed with a dose of 240 mJ/cm^2^. After flood exposure, substrates were developed using AZ726 MIF for 60 s. Development was stopped by rinsing with water. Exposed areas were etched using reactive ion etching with an RF power of 50 W, an inductively coupled plasma power of 100W, an SF6 flow rate of 20 sccm, an Ar flow rate of 10 sccm, and a processing time of 10 minutes (Oxford Plasmalab System 100). The sacrificial photoresist layer was removed by sonication in acetone and rinsing with isopropanol. The analysis of the electrophysiological data retrieved from the Multichannel System Software was performed employing a previously published algorithm for spike detection and sorting (wave_clus 3)^69^ using the programming and numeric computing platform MATLAB (R2021a).

#### Calcium Fluo-4 Assay

To verify the electrophysiological activity of hMOs on-chip, the intracellular calcium flux within the hMOs was analyzed employing a Fluo-4 Calcium Imaging Kit (Thermo Fisher) according to the manufacturer’s instructions (n=4). Videos were recorded using live-cell imaging (IX83, Olympus).

#### FITC Diffusion Study

5 kDa FITC-dextran (Thermo Fisher) was used for the assessment of dextran diffusion into hMOs on-chip (SI Figure 5A/B). To that end, a 100 μM solution of 5 kDa FITC-dextran in N2B27 medium was introduced into the media reservoirs of the microfluidic device. hMOs were exposed to the dextran solution for a period of 24 hours under both static and dynamic cultivation conditions, while transport into the microtissues was recorded simultaneously using live-cell imaging (IX83, Olympus). Influx and efflux into the organoids were analyzed by assessing the fluorescence intensity over time using the open-source image processing program FIJI.

### Image Analysis

#### Neurite Outgrowth Rate and Organoid Growth

Both maximum neurite outgrowth rate [μm/h] and organoid growth [μm^2^] were determined using the open-source image processing program FIJI. To determine the organoid growth, brightfield images of the hMOs were used to calculate the area of the individual organoids. To that end, images were transferred to 8-bit images; thresholds were adjusted to separate the hMOs from the background before converting the images to a mask and measuring the area using the measure function of the program. For the determination of the maximum neurite outgrowth rate, neurites extending from the hMOs were traced and measured using the freehand lines tool or the measure function, respectively.

#### Caspase 3

To analyze the apoptotic marker caspase 3 in the hMOs, images of the sectioned organoids were retrieved using a standard brightfield microscope (IX71, Olympus). The data analysis pipeline includes the adjustment of the contrast, color deconvolution, transfer to a binary image, generation of a mask, and measurement. The ratio of caspase 3 to total nuclei was subsequently calculated as follows:

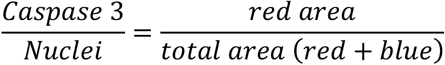

#### Immunofluore scence

The image analysis algorithms used in the study were applied as previously described.^70^

### Statistical Analysis

Statistical analysis and data visualization was conducted using the biostatistics program GraphPad Prism 8. For the assessment of statistical significance, Welch’s t-tests, Mann-Whitney tests, Kruskal-Wallis tests with Dunn’s multiple comparison tests, one-way, two-way ANOVAs or mixed-effect analysis with Geisser-Greenhouse correction and Tukey’s multiple comparison tests were performed. Normality was tested using a combination of the Shapiro-Wilk test and the Kolmogorov-Smirnov test. For the detection of outliers in normally distributed data sets, either a Grubb’s test for single outliers or a ROUST outlier test for multiple outliers was employed. Significances were assigned as follows: 0.12 (ns.), 0.033 (*), 0.002 (**), <0.001 (***).

## Acknowledgments

Graphical illustrations were generated using the online software Biorender. The study was supported by the Fonds National de la Recherche (FNR) Luxembourg (INTER/MERA/17/11760144, BRIDGES18/BM/12719664_MOTASYN). This work was supported by the U.S. Army Medical Research Materiel Command endorsed by the U.S. Army through the Parkinson’s Research Program Investigator-Initiated Research Award under Award No. W81XWH-17-PRP-IIRA. Opinions, interpretations, conclusions, and recommendations are those of the author and are not necessarily endorsed by the U.S. Army. We thank the Center for Micro- and Nanostructuring at the Technical University of Vienna for providing their clean room facilities.

## Author Contributions

SSp, PE, SB, and JCS conceived and designed the study. SSp and KB conducted microfluidic experiments. SSp, JF, and SS performed dopamine measurements. PS and HW fabricated MEA substrates. SSp and PS performed electrophysiological studies. TM provided expertise on oxygen sensing and costume-made microparticles for sensor integration. SSp, SB, BS, and JG performed immunohistochemical analysis. CJ and MH performed CFD simulations. SSp and SB conducted data analysis. All authors revised and approved the final manuscript.

## References

1. Maserejian, N., Vinikoor-Imler, L., Dilley, A. Estimation of the 2020 Global Population of Parkinson’s Disease (PD). in (MDS Virtual Congress 2020, 2020).

2. Poewe, W. et al. Parkinson disease. Nat. Rev. Dis. Prim. 3, 1–21 (2017).

3. Obeso, J. A. et al. Past, present, and future of Parkinson’s disease: A special essay on the 200th Anniversary of the Shaking Palsy. Mov. Disord. 32, 1264–1310 (2017).

4. Takahashi, K. & Yamanaka, S. Induction of Pluripotent Stem Cells from Mouse Embryonic and Adult Fibroblast Cultures by Defined Factors. Cell 126, 663–676 (2006).

5. Di Lullo, E. & Kriegstein, A. R. The use of brain organoids to investigate neural development and disease. Nat. Rev. Neurosci. 18, 573–584 (2017).

6. Cederquist, G. Y. et al. Specification of positional identity in forebrain organoids. Nat. Biotechnol. 37, 436–444 (2019).

7. Wang, Y., Wang, L., Zhu, Y. & Qin, J. Human brain organoid-on-a-chip to model prenatal nicotine exposure. Lab Chip 18, 851–860 (2018).

8. Monzel, A. S. et al. Machine learning-assisted neurotoxicity prediction in human midbrain organoids. Park. Relat. Disord. 75, 105–109 (2020).

9. Jo, J. et al. Midbrain-like Organoids from Human Pluripotent Stem Cells Contain Functional Dopaminergic and Neuromelanin-Producing Neurons. Cell Stem Cell 19, 248–257 (2016).

10. Tripathi, P. et al. Stem Cell Reports. Stem Cell Reports 9, 667–680 (2017).

11. Smits, L. M. et al. Modeling Parkinson’s disease in midbrain-like organoids. 1–8 (2019).

12. Nickels, S. L. et al. Reproducible generation of human midbrain organoids for in vitro modeling of Parkinson’s disease. Stem Cell Res. 46, (2020).

13. Abbott, N. J. Evidence for bulk flow of brain interstitial fluid: Significance for physiology and pathology. Neurochem. Int. 45, 545–552 (2004).

14. Mestre, H., Mori, Y. & Nedergaard, M. The Brain’s Glymphatic System: Current Controversies. Trends Neurosci. 43, 458–466 (2020).

15. Iliff, J. J. et al. A Paravascular Pathway Facilitates CSF Flow Through the Brain Parenchyma and the Clearance of Interstitial Solutes, Including Amyloid beta. 4, (2012).

16. Zou, W. et al. Blocking meningeal lymphatic drainage aggravates Parkinson’s disease-like pathology in mice overexpressing mutated α-synuclein. Transl. Neurodegener. 8, 1–17 (2019).

17. Whitesides, G. M. The origins and the future of microfluidics. Nature 442, 368–373 (2006).

18. Huh, D. A human breathing lung-on-a-chip. Ann. Am. Thorac. Soc. 12, S42–S44 (2015).

19. Campisi, M. et al. 3D self-organized microvascular model of the human blood-brain barrier with endothelial cells, pericytes and astrocytes. Biomaterials 180, 117–129 (2018).

20. Osaki, T., Uzel, S. G. M. & Kamm, R. D. Microphysiological 3D model of amyotrophic lateral sclerosis (ALS) from human iPS-derived muscle cells and optogenetic motor neurons. Sci. Adv. 4, 1–16 (2018).

21. Park, J. et al. Three-dimensional brain-on-a-chip with an interstitial level of flow and its application as an in vitro model of Alzheimer’s disease. Lab Chip 15, 141–150 (2015).

22. Rifes, P. et al. Publisher Correction: Modeling neural tube development by differentiation of human embryonic stem cells in a microfluidic WNT gradient (Nature Biotechnology, (2020), 38, 11, (1265–1273), 10.1038/s41587-020-0525-0). Nat. Biotechnol. 38, 1357 (2020).

23. Wang, Y., Wang, L., Guo, Y., Zhu, Y. & Qin, J. Engineering stem cell-derived 3D brain organoids in a perfusable organ-on-a-chip system. RSC Adv. 8, 1677–1685 (2018).

24. Cho, A. N. et al. Microfluidic device with brain extracellular matrix promotes structural and functional maturation of human brain organoids. Nat. Commun. 12, (2021).

25. Karzbrun, E., Kshirsagar, A., Cohen, S. R., Hanna, J. H. & Reiner, O. Human brain organoids on a chip reveal the physics of folding. Nat. Phys. 14, 515–522 (2018).

26. Ao, Z. et al. One-Stop Microfluidic Assembly of Human Brain Organoids to Model Prenatal Cannabis Exposure. Anal. Chem. 92, 4630–4638 (2020).

27. Yin, F., Zhu, Y., Wang, Y. & Qin, J. Engineering Brain Organoids to Probe Impaired Neurogenesis Induced by Cadmium. ACS Biomater. Sci. Eng. 4, 1908–1915 (2018).

28. Zhu, Y. et al. In situ generation of human brain organoids on a micropillar array. Lab Chip 17, 2941–2950 (2017).

29. Ray, L. A. & Heys, J. J. Fluid flow and mass transport in brain tissue. Fluids 4, (2019).

30. Zirath, H. et al. Every Breath You Take: Non-invasive Real-Time Oxygen Biosensing in Two-and Three-Dimensional Microfluidic Cell Models. Front. Physiol. 9, 1–12 (2018).

31. Zirath, H. et al. Bridging the academic-industrial gap: Application of an oxygen and pH sensor-integrated lab-on-a-chip in nanotoxicology. Lab Chip 21, 4237–4248 (2021).

32. Kieninger, J., Weltin, A., Flamm, H. & Urban, G. A. Microsensor systems for cell metabolism-from 2D culture to organ-on-chip. Lab Chip 18, 1274–1291 (2018).

33. Burré, J. The synaptic function of α-synuclein. J. Parkinsons. Dis. 5, 699–713 (2015).

34. Jensen, P. H., Nielsen, M. S., Jakes, R., Dotti, C. G. & Goedert, M. Binding of α-synuclein to brain vesicles is abolished by familial Parkinson’s disease mutation. J. Biol. Chem. 273, 26292–26294 (1998).

35. Ghiglieri, V., Calabrese, V. & Calabresi, P. Alpha-synuclein: From early synaptic dysfunction to neurodegeneration. Front. Neurol. 9, (2018).

36. Tsigelny, I. F. et al. Role of α-synuclein penetration into the membrane in the mechanisms of oligomer pore formation. FEBS J. 279, 1000–1013 (2012).

37. Lashuel, H. A., Overk, C. R., Oueslati, A. & Masliah, E. The many faces of α-synuclein: From structure and toxicity to therapeutic target. Nat. Rev. Neurosci. 14, 38–48 (2013).

38. Bousset, L. et al. Structural and functional characterization of two alpha-synuclein strains. Nat. Commun. 4, (2013).

39. Jarazo, J. et al. Parkinson’s Disease Phenotypes in Patient Neuronal Cultures and Brain Organoids Improved by 2-Hydroxypropyl-β-Cyclodextrin Treatment. Mov. Disord. 1–16 (2021) doi:10.1002/mds.28810.

40. Soofi, S., Last, J., Liliensiek, J., Nealy, P. & Murphy, C. The elastic modulus of MatrigelTM as determined by atomic force microscopy. 167, 216–219 (2009).

41. Leipzig, N. D. & Shoichet, M. S. The effect of substrate stiffness on adult neural stem cell behavior. Biomaterials 30, 6867–6878 (2009).

42. Hablitz, L. M. et al. Circadian control of brain glymphatic and lymphatic fluid flow. Nat. Commun. 11, (2020).

43. Tithof, J. et al. A network model of glymphatic flow under different experimentally-motivated parametric scenarios. iScience 25, (2022).

44. Modamio, J. et al. Synaptic decline precedes dopaminergic neuronal loss in human midbrain organoids harboring a triplication of the SNCA gene. bioRxiv 2021.07.15.452499 (2021).

45. Oueslati, A. Implication of Alpha-Synuclein Phosphorylation at S129 in Synucleinopathies: What Have We Learned in the Last Decade? J. Parkinsons. Dis. 6, 39–51 (2016).

46. Jo, J. et al. Lewy Body–like Inclusions in Human Midbrain Organoids Carrying Glucocerebrosidase and α-Synuclein Mutations. Ann. Neurol. 90, 490–505 (2021).

47. Jády, A. G. et al. Differentiation-Dependent Energy Production and Metabolite Utilization: A Comparative Study on Neural Stem Cells, Neurons, and Astrocytes. Stem Cells Dev. 25, 995–1005 (2016).

48. Takouda, J., Katada, S. & Nakashima, K. Emerging mechanisms underlying astrogenesis in the developing mammalian brain. Proc. Japan Acad. Ser. B Phys. Biol. Sci. 93, 386–398 (2017).

49. Wang, X. et al. Pathogenic alpha-synuclein aggregates preferentially bind to mitochondria and affect cellular respiration. Acta Neuropathol. Commun. 7, 41 (2019).

50. Paladini, C. A., Robinson, S., Morikawa, H., Williams, J. T. & Palmiter, R. D. Dopamine controls the firing pattern of dopamine neurons via a network feedback mechanism. Proc. Natl. Acad. Sci. U. S. A. 100, 2866–2871 (2003).

51. Stauffer, W. R. et al. Dopamine Neuron-Specific Optogenetic Stimulation in Rhesus Macaques. Cell 166, 1564–1571.e6 (2016).

52. Neufeld, T. P. Autophagy and cell growth - the yin and yang of nutrient responses. J. Cell Sci. 125, 2359–2368 (2012).

53. Lee, H. J. et al. Autophagic failure promotes the exocytosis and intercellular transfer of α-synuclein. Exp. Mol. Med. 45, 1–9 (2013).

54. Zagare, A., Gobin, M., Monzel, A. S. & Schwamborn, J. C. A robust protocol for the generation of human midbrain organoids. STAR Protoc. 2, (2021).

55. Song, J. J. et al. Cografting astrocytes improves cell therapeutic outcomes in a Parkinson’s disease model. J. Clin. Invest. 128, 463–482 (2018).

56. Berger, E. et al. Millifluidic culture improves human midbrain organoid vitality and differentiation. Lab Chip 18, 3172–3183 (2018).

57. Monzel, A. S. et al. Derivation of Human Midbrain-Specific Organoids from Neuroepithelial Stem Cells. Stem Cell Reports 9, 667–680 (2017).

58. Sackner-Bernstein, J. Estimates of Intracellular Dopamine in Parkinson’s Disease: A Systematic Review and Meta-Analysis. J. Parkinsons. Dis. 11, 1011–1018 (2021).

59. Klaentschi, K., Brown, J. A., Niblett, P. G., Shore, A. C. & Tooke, J. E. Pressure-permeability relationships in basement membrane: Effects of static and dynamic pressures. Am. J. Physiol. - Hear. Circ. Physiol. 274, 1327–1334 (1998).

60. Hsu, Y. H., Moya, M. L., Hughes, C. C. W., George, S. C. & Lee, A. P. A microfluidic platform for generating large-scale nearly identical human microphysiological vascularized tissue arrays. Lab Chip 13, 2990–2998 (2013).

61. Moreno-Arotzena, O., Meier, J. G., Amo, C. Del & García-Aznar, J. M. Characterization of fibrin and collagen gels for engineering wound healing models. Materials (Basel). 8, 1636–1651 (2015).

62. Ng, C. P. & Swartz, M. A. Fibroblast alignment under interstitial fluid flow using a novel 3-D tissue culture model. Am. J. Physiol. - Hear. Circ. Physiol. 284, 1771–1777 (2003).

63. Chee, P. N. & Pun, S. H. A perfusable 3D cell-matrix tissue culture chamber for in situ evaluation of nanoparticle vehicle penetration and transport. Biotechnol. Bioeng. 99, 1490–1501 (2008).

64. Madl, C. M. et al. Maintenance of neural progenitor cell stemness in 3D hydrogels requires matrix remodelling. Nat. Mater. 16, 1233–1242 (2017).

65. Reinhardt, P. et al. Derivation and Expansion Using Only Small Molecules of Human Neural Progenitors for Neurodegenerative Disease Modeling. PLoS One 8, (2013).

66. Njagi, J., Ispas, C. & Andreescu, S. Mixed ceria-based metal oxides biosensor for operation in oxygen restrictive environments. Anal. Chem. 80, 7266–7274 (2008).

67. Zanetti, C. et al. Monitoring the neurotransmitter release of human midbrain organoids using a redox cycling microsensor as a novel tool for personalized Parkinson’s disease modelling and drug screening. Analyst 146, 2358–2367 (2021).

68. Mika, J., Schwarz, K., Wanzenböck, H., Scholze, P. & Bertagnolli, E. Simultaneous Electrical Investigation of Isolated Neurites Using a Neurite-Isolation Device as Neurite Regeneration Model. in 9th Int. Meeting on Substrate-Integrated Microelectrode Arrays, 2014 vol. 9 322–323 (2014).

69. Chaure, F. J., Rey, H. G. & Quian Quiroga, R. A novel and fully automatic spikesorting implementation with variable number of features. J. Neurophysiol. 120, 1859–1871 (2018).

70. Bolognin, S. et al. 3D Cultures of Parkinson’s Disease-Specific Dopaminergic Neurons for High Content Phenotyping and Drug Testing. Adv. Sci. 6, 1–14 (2019).

