## Supplementary Information for "Development of a multi-sensor integrated midbrain organoid-on-a-chip platform for studying Parkinson’s disease"

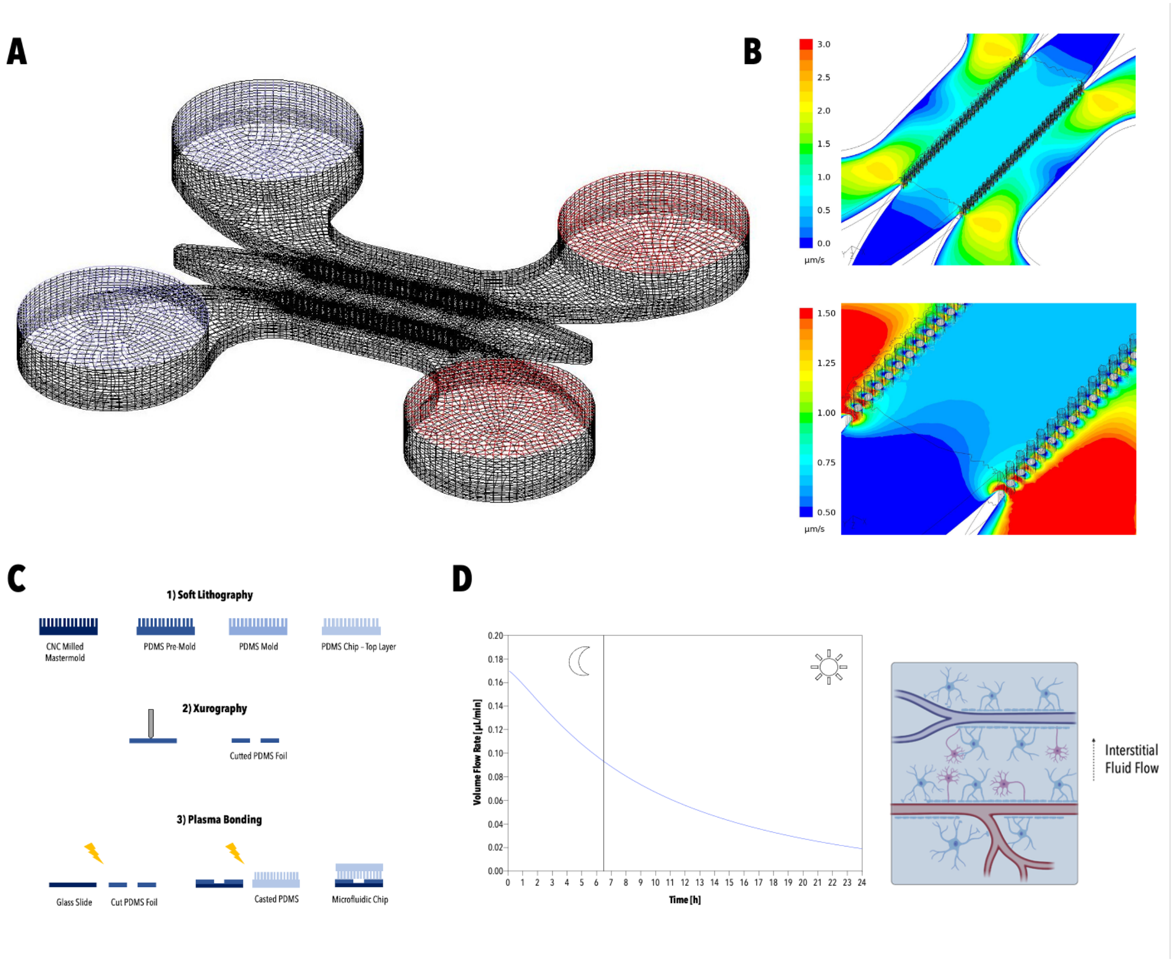


***SI Figure 1:*** *Graphical illustration of the hexahedral control volumes applied to the microfluidic device for CFD simulation (A). Overview of the simulated flow profile on chip depicting a highly uniform flow profile within the central hydrogel chamber as well as slightly higher flow rates at the corners of the hydrogel chamber (B). Workflow of the manufacturing process employed for the fabrication of PDMS devices (C). Time-resolved flow profile in the microfluidic device depicting circadian rhythms (D).*

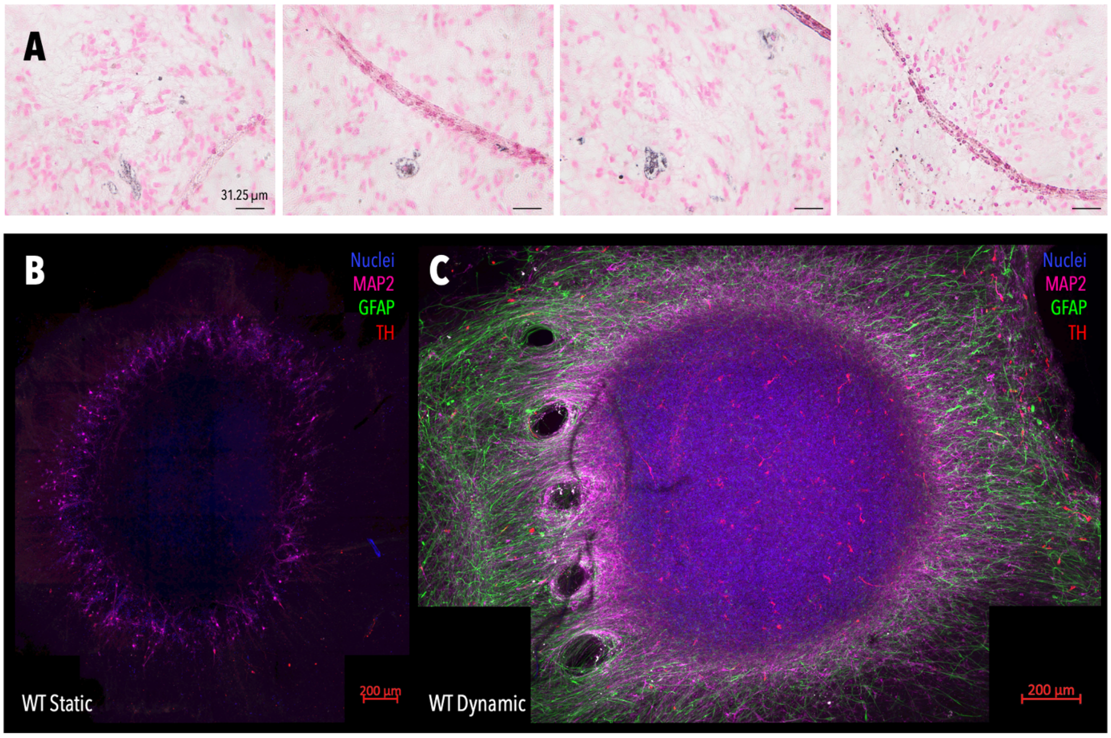


***SI Figure 2:*** *Fontana Masson staining of sectioned midbrain organoids revealing intra- and extracellular neuromelanin aggregates (A). Immunofluorescence staining of whole-mounted healthy hMOs cultivated under static culture conditions: TH (red), GFAP (green), MAP2 (magenta), nuclei (blue) (B). Immunofluorescence staining of whole-mounted healthy hMOs cultivated under dynamic culture conditions: TH (red), GFAP (green), MAP2 (magenta), nuclei (blue) (C).*

***SI Table 1:*** *Comparative analysis of immunohistochemically stained D60 whole-mounted hMOs cultivated under static or dynamic cultivation conditions respectively. Data was retrieved from three individual experiments.*

| | | #1 | #2 | #3 | Average | Dynamic/Static |
|---|---|---|---|---|---|---|
| Dynamic | TH/MAP2 [Px/Px] | 0.26116279 | 0.00420141 | 0.02983644 | 0.09840021 | |
| Static | TH/MAP2 [Px/Px] | 0.13835968 | 9.8965E-05 | 0.00260656 | 0.04702174 | **2.092653747** |
| Dynamic | GFAP [Px] | 0.26116279 | 0.00420141 | 0.02983644 | 0.09840021 | |
| Static | GFAP [Px] | 0.19976123 | 0.00215019 | 0.0162215 | 0.07271097 | **1.353306201** |

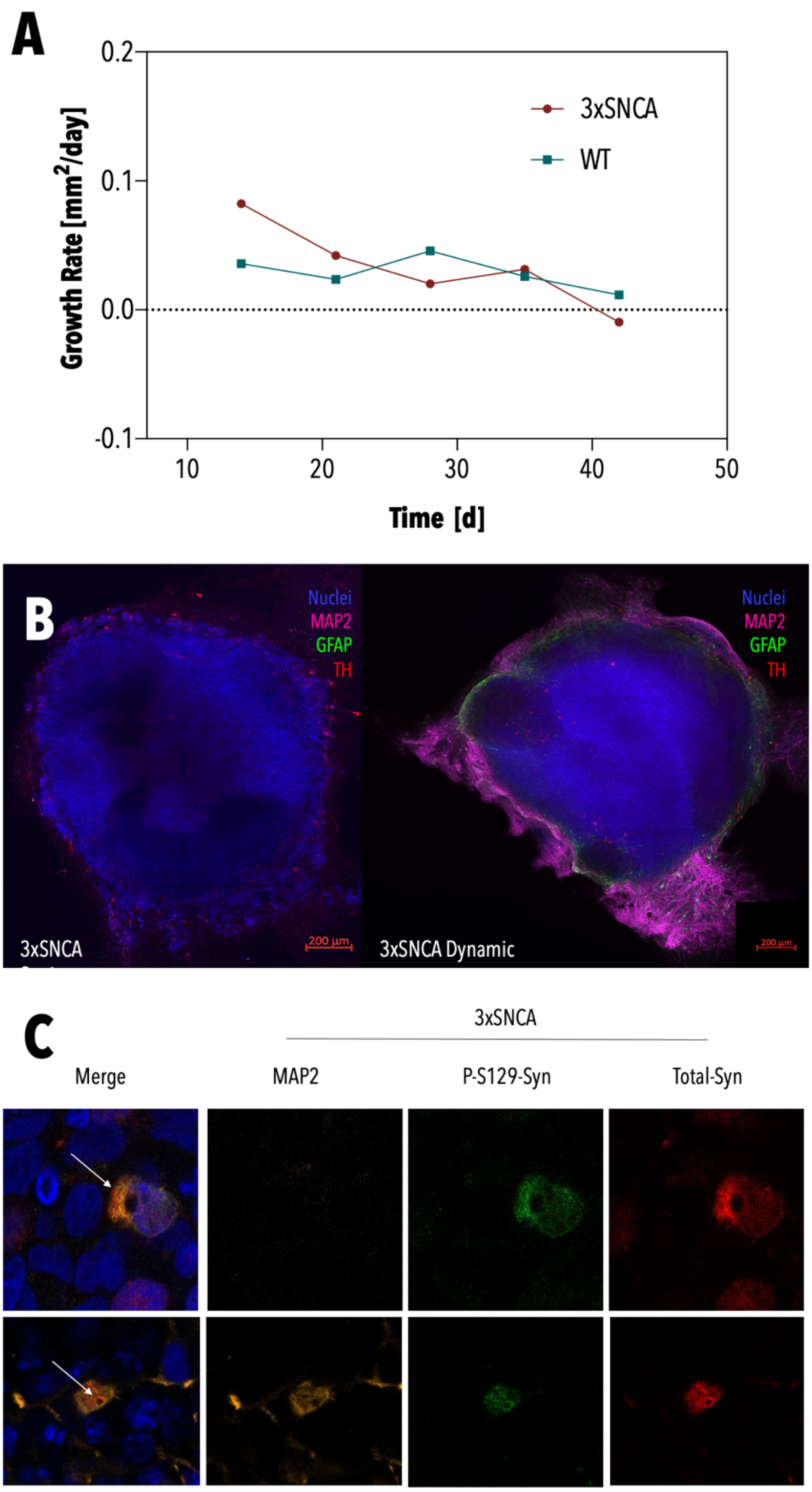


***SI Figure 3:*** *Average growth-rates of healthy and 3xSNCA hMOs (n=8-10 from 3 individual organoid batches) (A). Immunofluorescence staining of whole-mounted 3xSNCA hMOs cultivated under static culture conditions: TH (red), GFAP (green), MAP2 (magenta), nuclei (blue) (left panel). Immunofluorescence staining of whole-mounted 3xSNCA hMOs cultivated under dynamic culture conditions: TH (red), GFAP (green), MAP2 (magenta), nuclei (blue) (right panel) (B). Representative images of Lewy-body-like inclusions observed in 100 µm-thick sections of 3xSNCA hMOs (C).*

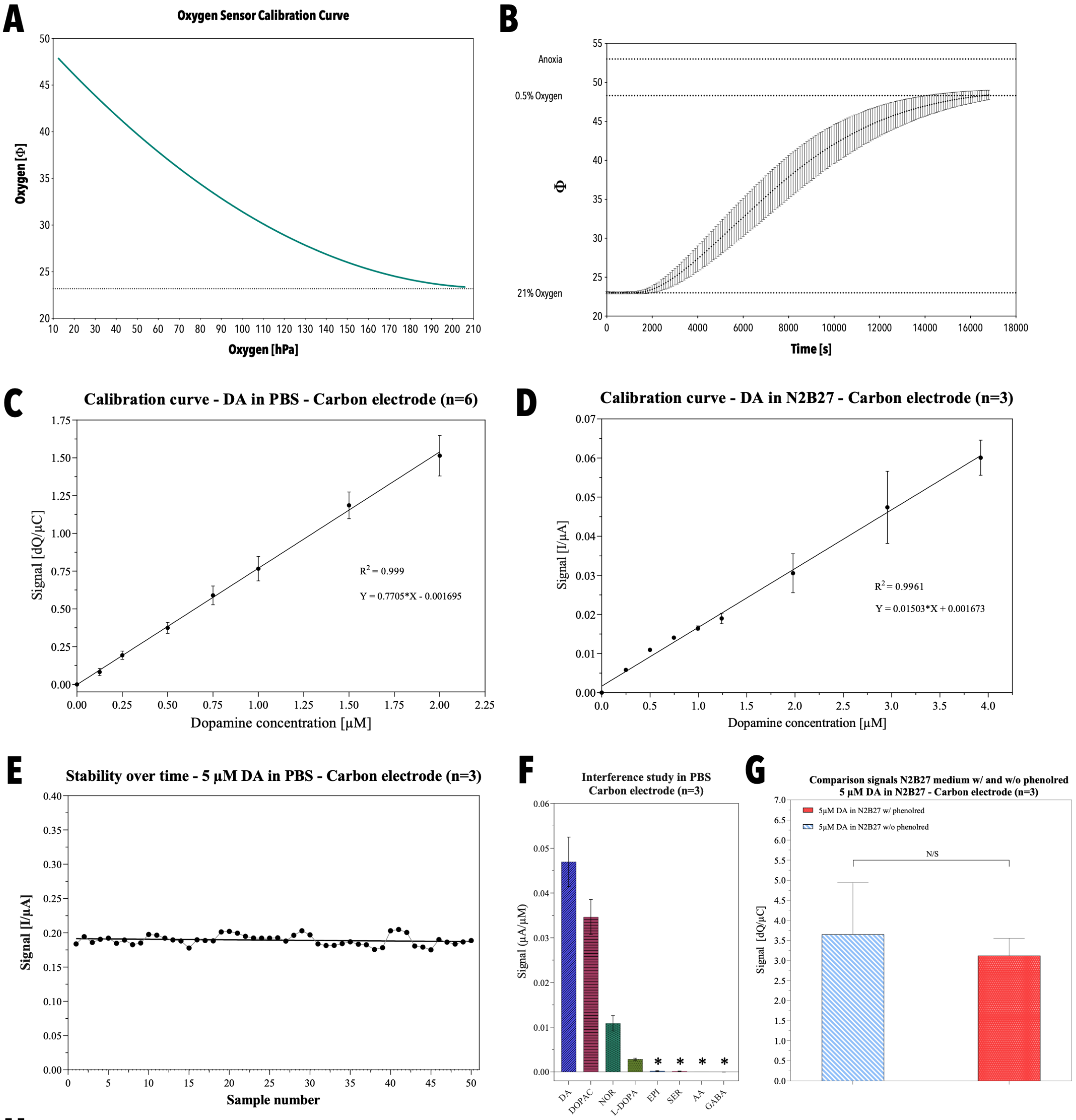


**H**

| | Average | SD | N |
|---|---|---|---|
| **LOD in PBS** | | | 6 |
| **LOD in Medium** | 145 nM | ± 0.7 nM | 4 |
| **CV** | % | | 7 |

***SI Figure 4**: Non-linear fit of oxygen sensor calibration curve on-chip (A). Recording of oxygen sensor calibration curve using an $O_2$-$CO_2$ controller (B). Dopamine calibration curve in PBS (n=6) and N2B27 (n=4) medium using a three-electrode setup (C/D). Graphical representation of sensor stability over 50 measurements at a dopamine concentration of 5 µM in PBS (n=3) (E). Interference study of common interferents in neurobiology: 3,4-dihydroxyphenylacetic acid (DOPAC), norepinephrine (NOR), levodopa (L-DOPA), epinephrine (EPI), serotonin (SER), ascorbic acid (AA), gamma-aminobutyric acid (GABA) (n=3). Studies were conducted in PBS. Significances were tested using a Welch's t-test *p<0.033, **p<0.002, ***p<0.001 (F). Comparative analysis of 5 µM DA signals retrieved in phenol red and phenol red-free cell culture media (n=3). Significances were tested using a Welch's t-test *p<0.033, **p<0.002, ***p<0.001 (G). Tabular overview of the DA sensor's sensitivity and reproducibility (H).*

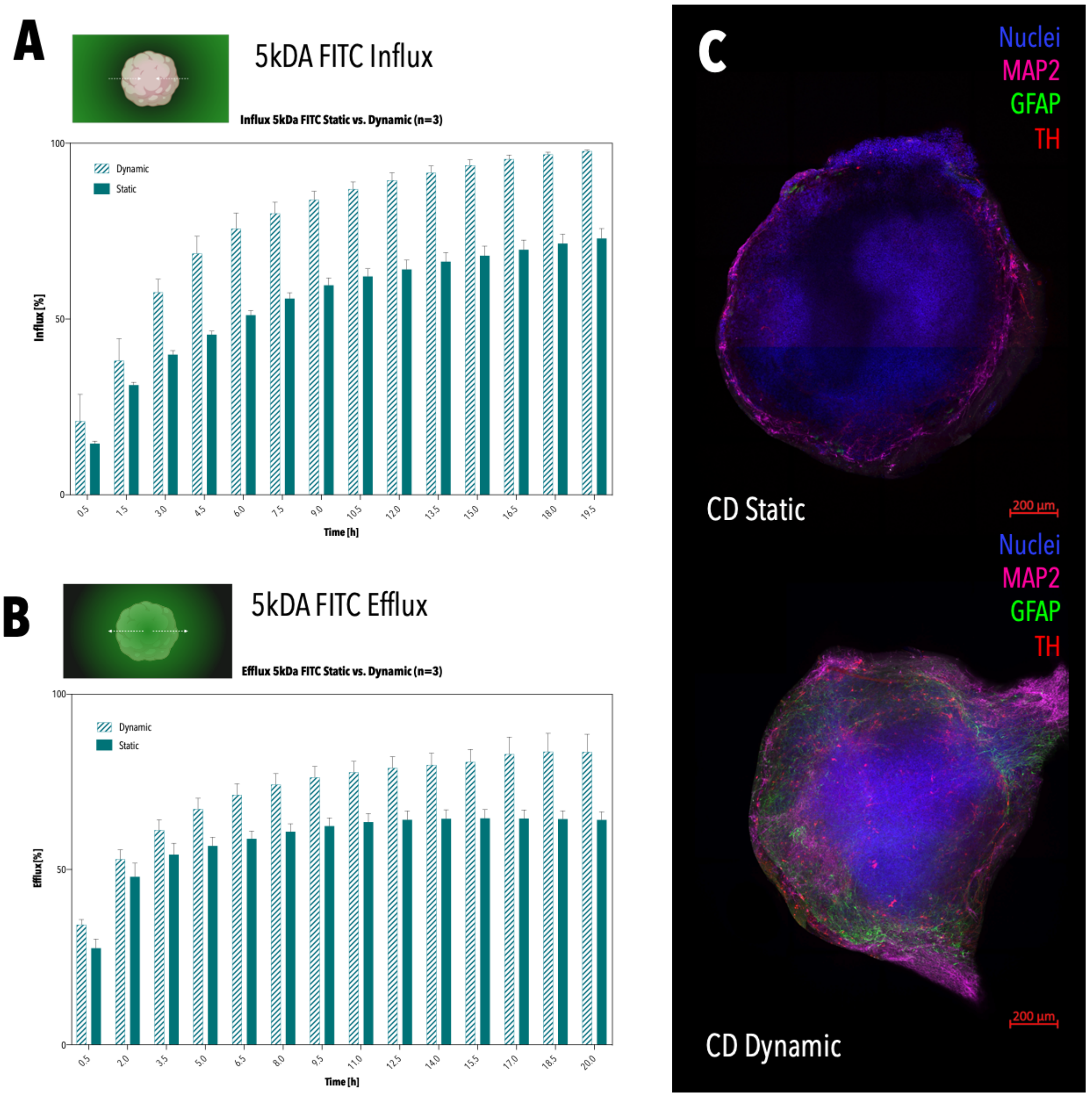


***SI Figure 5:*** *Time-resolved overview of 5kDA FITC transport into hMOs under static and dynamic culture conditions, depicting improved transport under dynamic conditions (n=3) (A). Time-resolved overview of 5kDA FITC transport out of hMOs under static and dynamic culture conditions, depicting enhanced removal of the dextran under dynamic conditions (n=3) (B). Immunofluorescence staining of whole-mounted 3xSNCA hMOs cultivated and treated with HP-β-CD under static culture conditions: TH (red), GFAP (green), MAP2 (magenta), nuclei (blue) (top panel). Immunofluorescence staining of whole-mounted 3xSNCA hMOs cultivated and treated with HP-β-CD under dynamic culture conditions: TH (red), GFAP (green), MAP2 (magenta), nuclei (blue) (bottom panel) (C).*

***SI Table 2:*** *Overview of used antibodies and respective dilutions.*

| **Target Protein** | **Dilution** | **Sample** | **Company** | **Catalog Number** | **RRID** |
|---|---|---|---|---|---|
| MAP2 | 1:200 | Whole-mount | Millipore | MAB3418 | AB_94856 |
| MAP2 | 1:1000 | Section | Abcam | ab92434 | AB_2138147 |
| GFAP | 1:200 | Whole-mount | Millipore | AB5541 | AB_177521 |
| TH | 1:200<br>1:1000 | Whole-mount<br>Section | Santa Cruz | sc-14007 | AB_671397 |
| TUJ1 | 1:1000 | Section | Millipore | AB9354 | AB_570918 |
| α-synuclein | 1:1000 | Section | Novus Biologicals | NBP1-05194 | n.a. |
| p-S129-α-synuclein | 1:500 | Section | Cell Signalling | 23706S | n.a. |
| TOM20 | 1:50 | Section | Santa Cruz | sc-17764 | AB_628381 |
